# Robustness and fidelity of voltage imaging analysis pipelines

**DOI:** 10.1101/2025.09.26.678573

**Authors:** Rui Silva, Hunor Bartalis-Szélyes, Daan Brinks, Zhenyu Gao

## Abstract

Tracking neuronal voltage changes using fluorescent voltage indicators is rapidly reshaping neuroscientific research. Voltage imaging enables direct visualization of electrical signals from subcellular compartments to large-scale networks, yet requires sophisticated image-analysis procedures. Here, we present a comprehensive study of current voltage imaging analysis pipelines and discuss the experimental conditions for which they are most suitable. We compare strengths and limitations of these pipelines in motion correction, denoising and segmentation routines, and discuss how different signal processing strategies can influence data integrity and interpretation. Our results show that most real-time analyses require GPU-accelerated algorithms and that denoising prior to signal processing is needed for analysis of subcellular dynamics. We conclude that voltage imaging analyses needs to be tailored to different experimental settings: we propose a decision-tree to determine analysis strategies for diverse experimental conditions. Together, these insights pave the way for reproducible, high-fidelity voltage imaging studies.

## Main

The importance of voltage imaging has been rapidly growing over the last decade. The increasing sensitivity, stability and brightness of genetically encoded voltage indicators (GEVIs)^1–3^, together with new imaging modalities and hardware^4–6^, have been paving the way for recordings with higher spatial and temporal resolution and signals with higher stability and signal-to-noise ratio (SNR). These rapid developments have led to a sprawl of voltage imaging studies using distinct voltage indicators, recording hardware, imaging modalities, illumination strategies and analysis pipelines (**Supplementary Fig 1a,b;** see **Supplementary Table 1** for all the references).

State-of-the-art voltage imaging allows a variety of experimental designs. Voltage imaging experiments can focus on specific neuronal populations from mesoscale^7–12^ to single cell resolution^13^ or even simultaneously record multiple populations in the same field-of-view (FOV)^2^. Moreover, the collected signals can range from slow oscillations to sub-millisecond electrical activity^2, 6, 8, 14, 15^, providing data to address in-depth questions about individual neurons, populations and neuronal networks. On a finer scale, several studies have also demonstrated the possibility to investigate voltage signals in subcellular compartments, such as dendrites^14, 16, 17^, axons^18–20^, dendritic spines^15, 21^ and even organelles, like the endoplasmic reticulum^22, 23^ and mitochondria^24^. Such high versatility and combination of high spatial and temporal resolution goes beyond the capabilities of current electrophysiology and calcium imaging techniques and demonstrates the suitability of voltage imaging to address questions that were previously challenging to tackle.

Signals obtained by voltage imaging vary in several key aspects. Voltage indicators differ in their brightness, kinetics, polarity and photostability. Their expression may vary between animals or cell types and their sensitivity is influenced by the illumination wavelength and imaging modality^25^. Over the years, reviews have discussed the developments in voltage indicators^26–31^, experimental methods and preparations^27, 32, 33^, or even new data that could be obtained with the progress in the field^34–36^. Less focus has been given on proper definition of analysis protocols for specific voltage imaging data. With the high experimental variability indicated above, tailored analyses pipelines are typically needed. Applying appropriate voltage imaging analysis requires an in-depth understanding of the specific experimental conditions, i.e. which imaging hardware, sensors and experimental protocols were chosen. Existing protocols are therefore typically not widely generalizable, and in-depth knowledge on their boundary conditions and limitations is needed to assess their suitability for analysing a particular experiment.

Voltage imaging analysis presents a number of challenges, namely motion artefacts, photobleaching, low SNR and other artefacts such as those induced by blood flow or camera shot-noise. Similar to calcium imaging, animal and tissue movements might introduce FOV shifts, especially during in vivo recordings, which can be corrected with standardized methods like NoRMCorr^37^. After correcting motion artefacts, a variety of denoising approaches can be applied to improve SNR, before or after segmentation of the regions of interest (ROI). Lastly, signal processing usually requires a baseline correction step, where unwanted fluctuations, such as those introduced by movement, blood flow, photobleaching and others are removed, followed by the extraction of fast electrical activity and/or slow oscillations (**Fig 1a**). Several analysis pipelines have been constructed to provide integrated solutions for these analytical challenges, with the intention of increasing standardization, improving data quality and promoting reproducibility.

**Figure 1.**
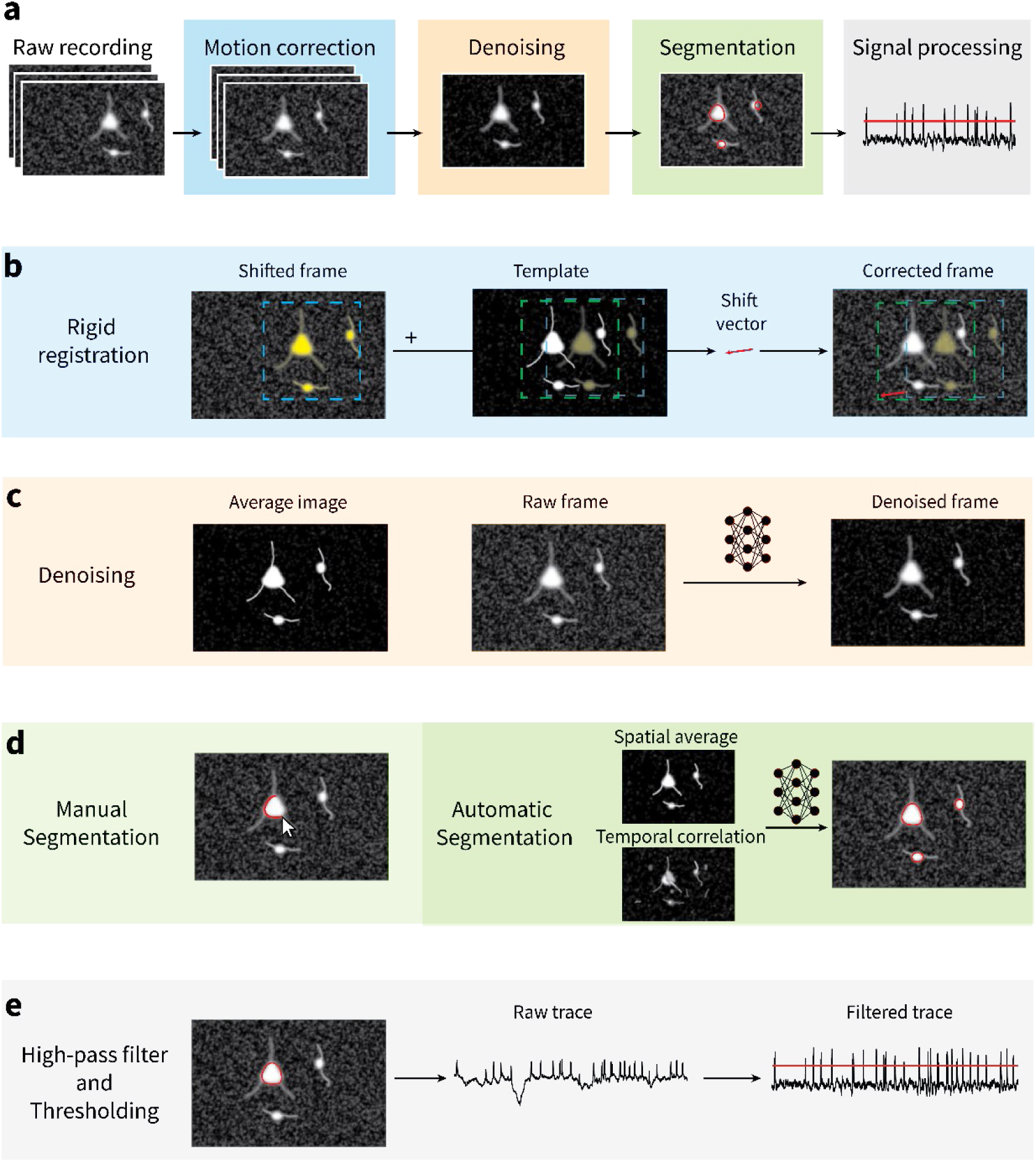
Representation of common voltage imaging analysis pipeline. **a)** Voltage imaging analysis steps. **b)** Representation of motion correction steps. Image is registered against a template, resulting in a shift vector applied to all pixels in a frame, thus correcting it. **c)** Representation of denoising pipeline aimed at individual pixel signal improvement, using neuronal network models. **d)** Representation of types of segmentation approaches, manual or automatic using neuronal network models. **e)** Representation of signal processing methods, based on correcting baseline fluctuations and spike detection.

In the context of the current capabilities and challenges of voltage imaging, here we performed a comprehensive analysis of several state-of-the-art analysis pipelines that are suitable for motion correction, denoising, segmentation and signal processing of voltage imaging signals. Since most published pipelines aim at accurate spike extraction from population neuronal voltage imaging with single cell resolution, we tested these pipelines on publicly available datasets with such characteristics. The datasets include data of different GEVIs (SomArchon, Voltron and paQuasAr-3s), recorded at different speeds and with widefield of or patterned illumination single-photon imaging. However, for each section we also discuss the most suitable methods for other data types, from mesoscale to sub-cellular resolution. For the first three steps: motion correction, denoising and segmentation, we perform an in-depth comparison of the more advanced and/or recent pipelines. For signal processing steps, we compared baseline correction methods for photobleaching and other artifacts, as well as spike detection approaches. We synthesize the insights from this analysis into a methodological flowchart, which will aid researchers in choosing and optimizing analysis pipelines tailored towards their specific experimental demands.

## Results

### State-of-the-art analysis pipelines for voltage imaging signals

To better understand the current analysis pipelines, we first performed a comprehensive search for recent voltage imaging papers and analysed the strategies researchers have been using to tackle the main artefacts present in voltage signals (**Supplementary Table 1**). We found that the analysis methods used by researchers are still highly variable (**Supplementary Fig 1c-f**). However, most pipelines included routines for motion correction (almost exclusively for *in vivo* studies), segmentation, spatial and/or temporal denoising and signal processing including baseline correction and spike detection (**Fig 1a**). The majority of motion correction strategies are based on registering individual frames towards a template, created by an average or median of several frames (**Fig 1b**). Denoising methods can aim at decreasing noise at a spatial or temporal level. Spatial denoising focuses on improving the signals of whole regions, by smoothing or averaging pixel traces, or on improving signals from individual pixels using signals from the neighbouring pixels and neighbouring frames^38–40^ (**Supplementary Fig 1di**). Besides improving SNR, spatial denoising can be useful to provide refined ROIs, since it can sharpen the neurons’ spatial footprint (**Fig 1c**). On the other hand, temporal denoising mainly focuses on removing unwanted fluctuations usually through frequency filters or removal of background fluctuations (**Supplementary Fig 1dii**). The segmentation was traditionally highly dependent on manual annotations (**Supplementary Fig 1e**), but some recent automatic pipelines could provide faster and more standardized results for somatic segmentation^41, 42^ (**Fig 1d**). Baseline correction is highly important prior to any other analysis of signals such as slow oscillations or spikes (**Fig 1e**), since an unstable baseline can prevent spike detection or falsely enhance the power of oscillations originating from artifacts.

In recent years a handful of published pipelines tackled almost all aforementioned steps: VolPy^41^, SGPMD-NMF^43^, FIOLA^44^, VOLTAGE^42^ and NOSA^45^, all of which are focused on electrical activity analysis. VolPy is based on motion correction, memory mapping, segmentation, photobleaching correction, spatial denoising and signal extraction^41^. FIOLA is a pipeline aiming at speeding up calcium and voltage imaging analysis through GPU computations, and presents similar routines to VolPy for voltage imaging analysis^44^. SGPMD-NMF has the same motion correction routine as VolPy, but is followed by different spatial denoising, segmentation and signal extraction pipelines^43^. VOLTAGE is focused on achieving reliable and fast segmentation, though GPU routines, but also has GPU accelerated motion correction with subsequent signal processing analysis^42^. NOSA is the only recently published pipeline with an interactive GUI for imaging analysis and it incorporates several routines for movement correction, background subtraction, baseline correction, segmentation, resampling and signal analysis^45^.

The aforementioned pipelines are not generalizable to all types of voltage imaging data, leading to researchers needing to adapt or develop their own analysis pipelines. However, some routines, as those used for motion correction or signal processing, can be used to analyse several types of data. To know which methods can be used for fast and accurate voltage imaging analysis, a thorough comparison of existing methods is needed. Thus, in the next sections we discuss existing methods, compare the best and/or more recent pipelines and suggest suitable methods for different types of data.

### Comparison of Motion Correction procedures

In recent years, in vivo voltage imaging with cellular or subcellular resolution has heightened the need for stable FOV recordings. However, motion artefacts, mostly from animal and brain movements, still persist in most recordings during behavioural experiments. Despite several studies opting for the removal of intervals or even full recordings where motion artefacts are present^7, 11, 46^, most researchers instead correct FOV shifts with available or customized methodologies (**Supplementary Table 1**). Some existing methods claim to perform automatic three-dimensional motion correction^47^, but as most voltage imaging recordings are still two-dimensional, this section focuses on 2D motion correction approaches.

We found that motion correction pipelines are relatively consistent among papers (**Supplementary Fig 1c**). Around 40% of the analysed papers use NoRMCorr with rigid transformations^37^ or derived methodologies, while another 17% used other customised rigid transformations. NoRMCorr is based on matching frames to a template through maximisation of the normalised cross correlation (NCC) and this method has been integrated in the CaImAn package^48^ and in published pipelines^41, 43^. Several studies also used routines from the Fiji software, namely TurboReg, which is based on an old mean-squared-error (MSE)^49^ method with further image warp, and TrackMate^50^, whose particle tracking routine can be used to infer image shifts^51^. However, while NoRMCorr routine was already adapted for batch processing to previously built templates^52^, software such as Fiji requires loading the complete dataset into memory, hindering analysis of heavy datasets on weaker workstations. These CPU-based routines are slow and computationally heavy, and with the increasingly large datasets that can be recorded, researchers would benefit from faster pipelines. Recently, two pipelines, FIOLA and VOLTAGE, incorporated GPU-accelerated routines for motion correction, based on similar normalized cross correlation (NCC) principles to that of NoRMCorr^42, 44^, that could tackle this problem.

Here, we compared the state-of-the-art NoRMCorr pipeline, with the faster FIOLA and VOLTAGE pipelines. NoRMCorr has mainly two parameters that heavily influence the motion correction output: the maximum shifts in x and y coordinates that a frame can have versus the template; and the size of a gaussian filter (gSig) to denoise the images prior to calculate the shifts. We found that the shifts calculated with and without a gaussian filter can differ substantially (**Fig 2a**), with the motion correction being significantly better with application of the filter. As FIOLA GPU motion correction performs similarly to NoRMCorr without filter, we added the same filter to the FIOLA pipeline and verified significant changes to the shifts calculation, similarly to those observed using NoRMCorr (**Fig 2b**). While the shift differences between filtered and unfiltered data were almost negligible for most recordings, in some cases the errors increased up to 10 pixels (**Fig 2c**), which can significantly change the resulting traces of recorded cells. We noticed the largest differences occurring for recordings where the cell to background contrast was very low (**Fig 2ei)**. It is important to note that this gaussian filter uses a vignette kernel, rather than a normal gaussian kernel (**Fig 2d**), resulting in improved cell edges, which allow for better shift calculation (**Fig 2eii-2eiii**).

**Figure 2.**
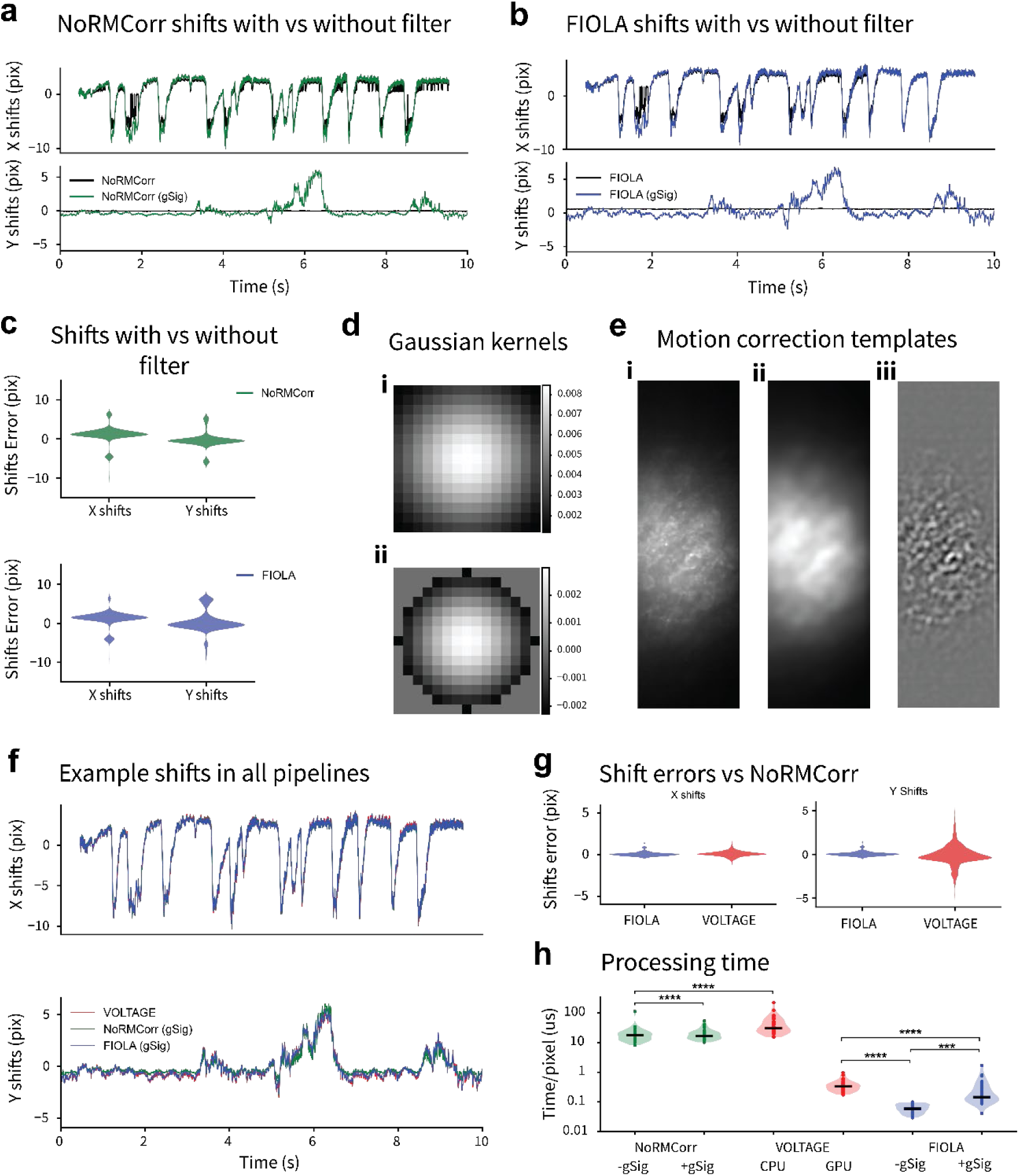
Comparison of motion correction pipelines. **a)** Example shifts calculated with NoRMCorr with (green) and without (black) vignette gaussian denoising. **b)** Example shifts calculated with FIOLA with (green) and without (black) vignette gaussian (gSig) blur. **c)** Shifts difference of computations with versus without vignette gaussian blur computed by NoRMCorr (green) and FIOLA (blue). **d)** Gaussian kernels used to compute panels eii (**i**) and eiii (**ii**). **e)** Example templates used for motion correction. **i)** Average image of the first 1000 frames of the recording. **ii)** Gaussian blur of the average image of panel **i)**. **iii)** Vignette gaussian blur of the average image of panel **i)**. **f)** Example shifts calculated with VOLTAGE (red), NoRMCorr (green) and FIOLA (blue) with vignette blur. **g)** Shifts difference between FIOLA or VOLTAGE versus NoRMCorr. **h)** Timing per pixel to compute motion correction with the different pipelines (n=37 videos). All panels in this figure correspond to the analysis of the representative recording 00_03 of the HPC2 dataset, which was shown as an example for having one of the biggest shifts calculated among all videos of all 4 datasets. Panels C and G only compute the differences between frames where shift computation in NoRMCorr was bigger than 4 pixels (n=1570). X shifts correspond to shifts in image width axis. Y shifts correspond to shifts in image height axis. gSig corresponds to the computation of vignette gaussian blur. Statistical analysis was computed as detailed in the methods section, and the significance shown corresponds to post-hoc tests.

VOLTAGE solely performs an intensity normalisation prior to motion correction calculation, and instead of calculating NCC on the whole image, it calculates it in patches, providing a quite accurate shift estimation (**Fig 2f**). Both FIOLA and VOLTAGE perform comparably to NoRMCorr, with FIOLA showing lower shift differences (<2.5pix) than VOLTAGE (<5pix) (**Fig 2g**). However, on our setup, FIOLA processing speed was significantly faster than VOLTAGE, similarly to NoRMCorr being faster than VOLTAGE processed on the CPU (**Fig 2h**). Since VOLTAGE only works on Linux operating system (OS), and we tested it in Windows Subsystem for Linux, these results might differ on a native Linux computer.

These results show that both VOLTAGE and FIOLA can perform comparably to the state-of-the-art NoRMCorr with more than 10x faster processing speed. However, FIOLA requires the addition of the gaussian filter from NoRMCorr for accurate correction and VOLTAGE is slower than FIOLA. Moreover, while FIOLA can be used on any OS and correct videos with previously calculated templates, VOLTAGE can only be used on Linux and only corrects the frames to a template created with the loaded frames. Thus, the FIOLA motion correction pipeline is more versatile and suitable for future uses in pipelines aiming at real-time analysis.

### Comparison of denoising procedures

Voltage imaging recordings can have low SNR for several reasons. The use of membrane localised indicators with low brightness and/or low sensitivity, out-of-focus fluorescence crosstalk between neighbouring cells, and recording artefacts, such as those induced by motion, blood flow or recording systems, can influence the images collected during recordings. Denoising methods can focus on spatial or temporal corrections, depending on the targets of study and the necessary spatial-temporal resolution.

The choice of temporal denoising methods is primarily determined by the required temporal resolution of specific experiments. Slow oscillation studies usually use simple methods such as frequency filters or multi-trial signal averaging (**Supplementary Fig 1eii; Supplementary Table 1**). For such studies, if heartbeat frequency contaminates signals of interest, it is usually filtered out by band-pass filters^9, 53–56^. On the other hand, such strict methods are usually not suitable for studies focused on fast electrical activities such as action potentials, as the temporal resolution is of utmost importance. We found that regardless of the target of study, dark current and background subtraction are commonly applied (**Supplementary Fig 1eii**). The dark current refers to the camera noise before illuminating the sample of interest; and the background refers to an average value obtained from a region of the sample where cellular fluorescence is not seen or expected. While subtracting dark noise can only provide more accurate ΔF/F values, subtracting background can remove noisy fluctuations, thus improving the voltage signal quality. Several studies select background regions surrounding the cells of interest^51, 57, 58^, while others arbitrarily performed manual annotations or use all the non-labelled region as the background for subtraction^16, 59^. However, caution is necessary when selecting background regions, as inappropriate choices may inadequately remove noise fluctuations or even introduce artifacts. For example, incorporating areas with blood flow oscillations or out of focus signals, can substantially compromise the signal quality. Consequently, particular care must be taken when analysing signals sensitive to background contamination, especially for in vivo imaging studies.

Spatial denoising methods also differ with the demand of spatial resolution. In mesoscale studies, simple methods such as gaussian or average spatial filters are the first choices (Supplementary Fig 1eii**; Supplementary Table 1**). In contrast, for cellular and subcellular voltage imaging, variations across different pixels may be valuable. Thus, the best denoising methods should focus on improving the signals of each individual pixel. Denoising individual pixels in low SNR recordings is quite challenging, thus it is no surprise that the methodologies used are quite variable (**Supplementary Table 1**). Some studies used approaches such as penalized matrix decomposition (PMD)^15, 58, 60^ or Principal Component Analysis (PCA)^61, 62^, while others used approaches that intertwine segmentation and denoising to select or enhance pixels with more information, such as such as the case of PCA-ICA^63^ or pixel weighting algorithms (more details in the *Segmentation* section). Published pipelines have incorporated some of these strategies^41, 43^. However, these methods are time-consuming and not always improve voltage signals. Emerging new denoising pipelines based on self-supervised learning methods may provide a better denoising performance. The DeepVID pipeline denoises each frame using a sequence of frames surrounding it^64^, and the further development DeepVIDv2, improves the initial framework by adding an edge extraction method to enhance the denoising of small structures^38^. The CellMincer pipeline also takes into account spatial features in the image, which could substantially improves the denoising accuracy^40^. The SUPPORT pipeline uses a spatio-temporal algorithm where the influence of neighbouring pixels in the central frame is weighed stronger in the denoising process than the temporal characteristics of the signal in the pixel of interest^39^. It is suggested that these methods are superior to PMD and other self-supervised pipelines developed for denoising calcium imaging data, such as DeepCAD-RT, DeepInterpolation and Noise2Self. Therefore, a side-by-side comparison of their specific training requirements and performance would be informative.

We analysed the training procedures and denoising performance of DeepVIDv2, SUPPORT and CellMincer, using publicly available datasets. While DeepVIDv2 and SUPPORT require one line of code with user selected parameters and datasets to train a model, CellMincer requires metadata files for data specification, preprocessing and training. Moreover, DeepVIDv2 and SUPPORT denoising inference requires recordings and a model, whereas CellMincer requires a CPU based preprocessing step before denoising with the trained model, which is slightly less user-friendly. These pipelines also differ in their requirements for training datasets. For SUPPORT and CellMincer a training dataset could consist of a single video of 3000 and 5000 frames, respectively, to achieve satisfactory results. On the other hand, DeepVIDv2 used 1000 videos of 1000 frames, and DeepVID^64^ used 1181 videos of 1000 frames for training. In addition to this requirement for a large training dataset, combining videos with different height and width into the same training dataset is also challenging. Based on these constraints, SUPPORT was easiest to implement.

Since the DeepVIDv2 pipeline assumes the dataset is free of motion artefacts^38^, for a fair comparison, we trained all models with the same dataset after correction of motion artefacts using NoRMCorr (for more information see *Methods*), and further denoised the training dataset for initial analysis. Upon visual inspection, we noticed the successful removal of zero-mean gaussian noise from the recordings by all three pipelines, leading to improved visibility and localization of the cell’s activity (**Fig 3a**). However, while the spikes waveforms after SUPPORT and CellMincer processing were largely retained, the spikes waveforms after DeepVIDv2 processing became smaller and broader (**Fig 3b**). Moreover, all pipelines decreased the SNR of the whole cell mask trace (**Fig 3c**). We postulated that this could be a result of overfitting or undertraining. However, the average Normalized Cross Correlation (NCC) between each frame and the average z-stack projection was hardly affected with increasing iterations of training (**Supplementary Figure 2**). Moreover, the spike waveforms and SNR remained largely constant after 2 iterations of DeepVIDv2, 20 iterations of SUPPORT and 2000 iterations of CellMincer (**Supplementary Figure 3; Fig 3c**). On the other hand, all pipelines improve the average SNR of signals of individual pixels (**Fig 3d**). These results suggest that the pipelines were successful at removing zero-mean gaussian noise from each individual pixel, improving their signals. However, the result of this denoising process is not significantly better than averaging a large number of pixels of ROIs with the same signals

**Figure 3.**
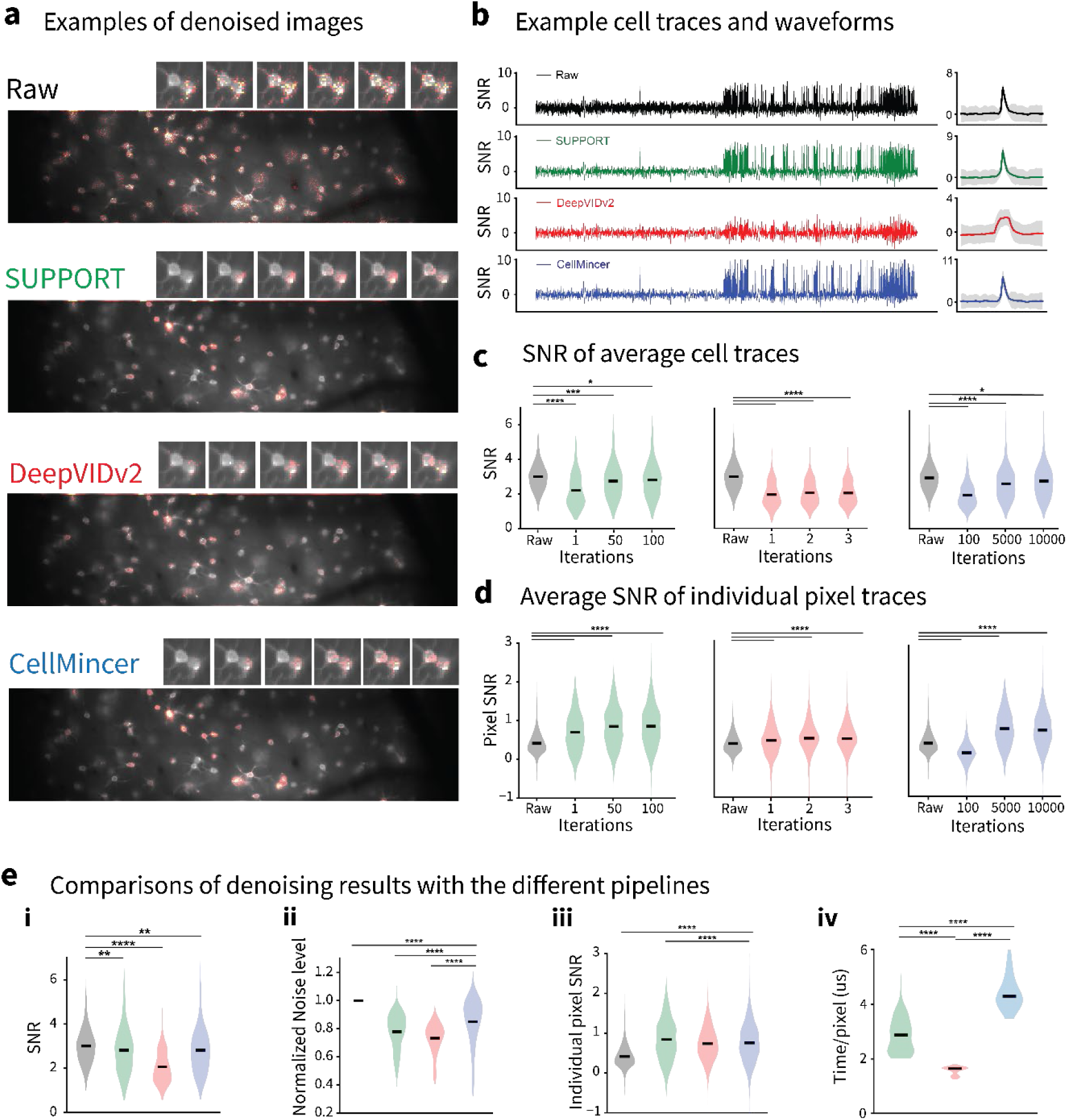
Comparison of denoising pipelines. **a)** Examples of raw and denoised images with different pipelines. **b)** Example cell traces (left) and waveforms (right), obtained by averaging all pixels of the ground-truth cell masks. **c)** Average spike SNR of cell traces of L1 dataset with SUPPORT (green), DeepVIDv2 (red) and CellMincer (blue), with models obtained with different training iterations **d)** Average SNR of all individual pixels from the ground-truth masks of one recording. **e)** Comparison of denoising results with 100 iterations model of SUPPORT (green), 3 iterations model of DeepVIDv2 (red) and 10000 iterations model of CellMincer (blue). **i)** Average spike SNR of cell traces of L1 dataset. **ii)** Normalised noise level of cells’ traces. **iii)** Average single pixel SNR. **iv)** Computation time per pixel of each pipeline in all videos of L1 dataset. The datapoints of panels C, Ei and Eii are the average of each neuron in the L1 dataset, with n=483. The datapoints of panels D and Eiii are the average of individual pixels of the ground-truth masks of the recording L1.00.00 of L1 dataset, with n=7235. Statistical analysis was computed as detailed in the methods section, and the significance shown corresponds to post-hoc tests.

Cross comparison of their overall performance showed that the DeepVIDv2 decreases the SNR substantially more than SUPPORT and CellMincer (**Fig 3ei**). This is accompanied by a reduction of the normalized noise level, suggesting that the denoising process removes both noise and signal (**Fig 3eii**), consequently distorting spike waveforms (**Fig 3b**). All the pipelines were successful in improving the average pixel SNR, with SUPPORT having the best performance among all three (**Fig 3eiii**). Given that all studies reported that their models can be generalised and applied to other recordings, we also denoised other datasets using the same trained models. We analysed both HPC and HPC2 datasets, which differ not only in the polarity of the voltage indicator, but also in the illumination strategy. Out of the three pipelines, SUPPORT showed the best results, by better conserving the spike waveform (**Supplementary Figure 4a,b**) and for providing the smallest decrease in the trace SNR, together with the higher single pixel SNR increase (**Supplementary Figure 4c-d**). Lastly, by comparing the pipelines processing speed, we could conclude that all of them are quite slow, allowing for analysis of less than 1000 pixels per millisecond, with DeepVIDv2 being the fastest, and CellMincer being the slowest (**Fig 3eiv**).

Overall, these pipelines showed clear removal of zero-mean gaussian noise. However, the improvement of the voltage signals might be accompanied by a change of spike waveforms. Moreover, the slow processing speed might hinder their application for analysis of big datasets aiming at fast results collection. The SUPPORT and CellMincer pipelines are effective in improving the SNR of individual pixels, which could be suitable for the analysis of sub-cellular signals.

### Comparison of Segmentation procedures

Signal extraction requires the selection of the regions of interest (ROI), which, depending on the research question, can vary from broad areas (such as mesoscopic functional regions), single neurons, to subcellular compartments. For studies focusing on mesoscale analysis, manual annotations are frequently required, as the ROIs vary between studies and the FOV recorded can range from small regions of an ex-vivo slice^65^ to large cortical areas^53, 56^. Moreover, since mesoscale imaging is typically done with dense expression of voltage indicators and focuses on regional fluctuations, automatic segmentation based on either fluorescence values or activity is challenging. Likewise, segmentation of subcellular compartments or organelles can also benefit from manual segmentation. Defining portions of axons, dendrites and organelles may require dividing a recorded cell into subcellular regions that present very small amplitude signals^15, 16, 22–24, 66, 67^. These constraints make the current automatic methods insufficient for a proper segmentation of specific ROIs.

In contrast, voltage imaging focusing on well isolated cells can benefit from the accuracy and speed that automatic segmentation can provide. Currently, automatic segmentation methods are still not standardised, but several strategies were applied to automatically segment signals. The most common automated pipeline we found is based on the PCA/ICA analysis originally applied to calcium imaging^63^. A few papers also used a semi-automated maximum-likelihood pixel-weighting (MLPW) algorithm, which weights pixels of an initially given mask based on their correlation with the signal of the whole mask^17, 68, 69^. In addition, despite the accuracy problems that heterogenous illumination or brightness might bring, several papers still applied intensity-based thresholding method for segmentation (**Supplementary Table 1**). Pipelines such as these can be inaccurate, to segment ROIs with low amplitude signals, and can be slow and computationally heavy, which defies the purpose of automatic segmentation. To circumvent these issues, two recent studies implemented neuronal network based models, namely Mask R-CNN^41^ and VOLTAGE^42^to achieve faster and more accurate segmentation of somatic ROIs.

To evaluate the speed and accuracy of these two voltage imaging segmentation pipelines, we analysed the performance of VOLTAGE and Mask R-CNN on several publicly available voltage imaging datasets (**Fig 4a**). Both pipelines achieved high F1-scores for different datasets, with the exception of Mask R-CNN for the HPC2 dataset, with values in alignment with what was previously reported^41, 42^ (**Fig 3b, Supplementary Fig 5a**). Consistent with the claim from the original study^42^, VOLTAGE was significantly faster than Mask R-CNN across datasets (**Fig 4c, Supplementary Fig 5b**). This is largely due to a significant decrease in the time taken for the calculus of summary images, as the model prediction stage for Mask R-CNN is still faster than that of VOLTAGE (**Fig 4c, Supplementary Fig 5b**). The summary images of both pipelines are based on spatial averages and temporal correlations, whose computation can be dependent on the number of frames used. For this reason, we interrogated how both pipelines performed under an increasing number of frames and what would be the minimum number of frames required for optimal performance. We tested the segmentation accuracy and speed of both approaches using the first 1000, 2000, 5000, 10000 frames and the complete videos. Both VOLTAGE and Mask R-CNN had robust and high accuracy performance from just 1000 frames in the L1 dataset (**Fig 4d**), but also in the TEG and HPC datasets (**Supplementary Fig 5c)**. However, only VOLTAGE showed significant accuracy improvements with increasing video lengths in the HPC2 dataset (**Fig 4d**). Lastly, we also found that VOLTAGE is faster than Mask R-CNN across all video lengths and datasets (**Fig 4e; Supplementary Fig 5c**).

**Figure 4.**
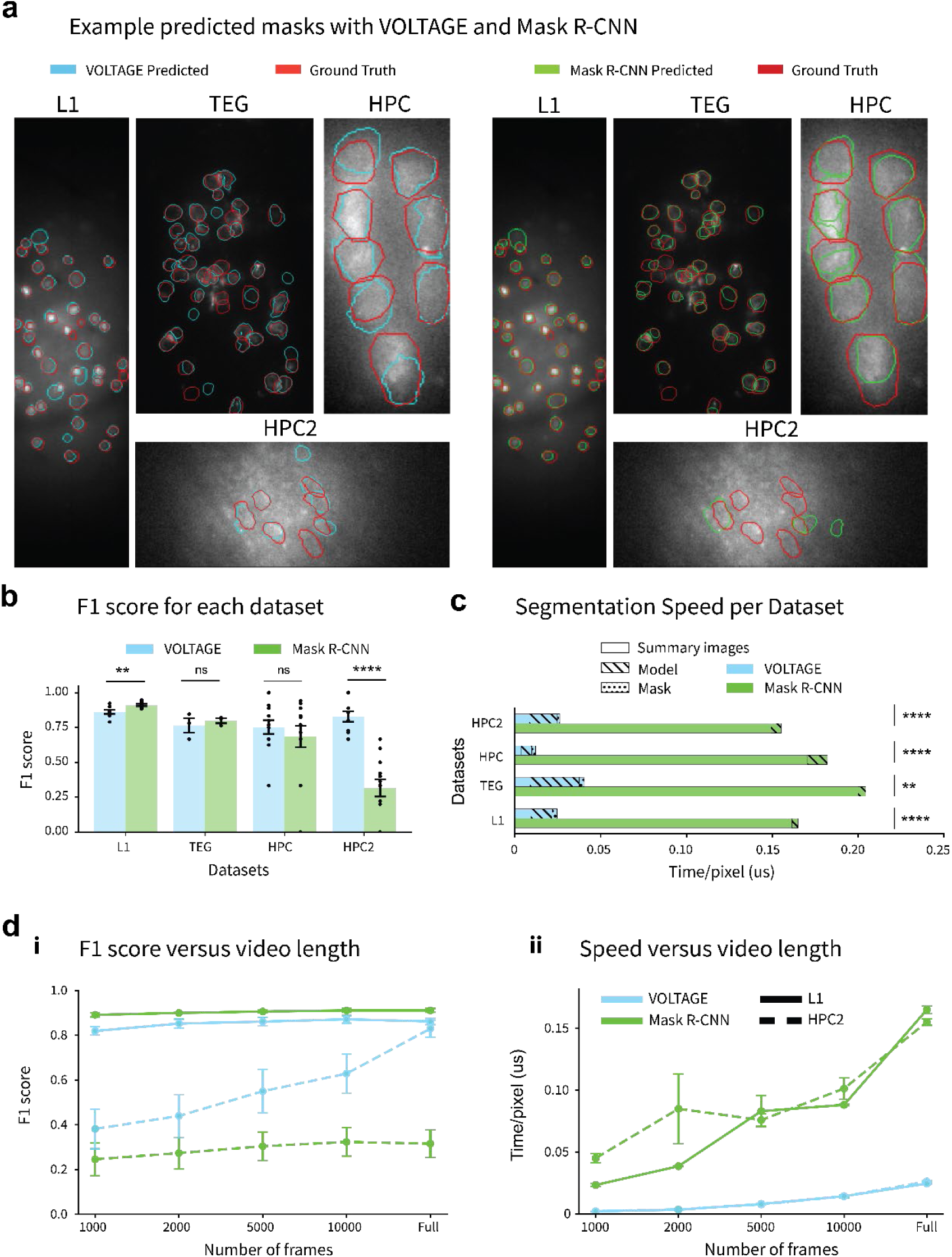
Comparison of segmentation pipelines. **a)** Example of ground-truth (red) and predicted masks with VOLTAGE (blue) and Mask R-CNN (green), in different datasets. **b)** F1 scores per dataset of VOLTAGE and Mask R-CNN. **c)** Average segmentation processing speed in each dataset, divided per segmentation steps. **d)** F1 scores (**i**) and speed (**ii**) of each segmentation pipeline, in the L1 and HPC2 datasets, with increasing number of frames. Each video cropping starts on the first recorded frame. The number of data points for each graph are the number of videos of the dataset: L1 - 9 videos; TEG 3 videos; HPC 12 videos; HPC2 - 13 videos. Statistical analysis was computed as detailed in the methods section.

The low initial performance of both pipelines on the HPC2 dataset might be a result of its low SNR and/or the low cell-to-background fluorescence contrast (**Fig 4a**). However, the unique increase in accuracy of VOLTAGE with increasing number of frames suggests that Mask R-CNN performance might be independent on the number of frames above 1000 frames, while for low contrast videos VOLTAGE can still be improved with increasing video length. In conclusion, our results show that, while both VOLTAGE and Mask R-CNN attain similarly high levels of accuracy in several datasets, VOLTAGE achieves faster processing speed and seems to be able to improve its performance with increasing video lengths.

### Comparison of signal processing procedures

The performance of GEVIs have been greatly improved over the years, allowing for longer recordings of large population of cells. However, longer recordings are more susceptible to fluorescence decay due to photobleaching. Such gradual fluorescence decay could affect subsequent spike detection and accurate analysis of oscillation. The photobleaching can be partially mitigated by using lower light power and by proper signal processing of voltage traces. However, defining the best methodology to use can be quite challenging, since the degree of fluorescence decay can vary greatly, depending on the specific indicator, light power, expression density and illumination strategy (**Supplementary Fig 6a**). Besides this, voltage signals can also have baseline fluctuations due to artefacts such as brain movement or blood flow, even when photobleaching effects are insignificant (**Supplementary Fig 6b**), or present plateau like signals during bursts of activity (**Supplementary Fig 6a, red squares**). Separating all fluctuation sources in independent problems is very challenging, thus baseline correction routines should tackle all present instabilities together.

Several processes such as exponential fits, high-pass filters and rolling averages/medians for baseline corrections are commonly applied for baseline corrections (**Supplementary table 1, Supplementary Figure 1f)**. Subtracting an exponential fit to the signal trace seems a natural choice for correcting photobleaching, especially since voltage sensors are membrane localized and have lower diffusion constants than cytosolic fluorophores. However, this is only suited for close-to-ideal measurements with high sensor fluorescence-over-background, high SNR of spikes, and photobleaching being the dominant noise source, despite have low amplitude components in the frequency range of interest of electrical activity. (**Supplementary Fig 6ci**). Most measurements do not fulfil this stringent condition, and exponential fits are therefore not generally suitable for baseline correction (**Supplementary Fig 6cii**). Frequency filters, mainly high-pass, and rolling functions, such as average or median, can output similar results for similar analysis windows (**Supplementary Fig 6ciii-vi**). High-pass filters remove all frequencies up to a certain cut-off value, while a rolling average removes the average of a window around data points. As an example, when using 1/3Hz high-pass filters, all slow oscillations from 3 seconds upwards are removed, creating a similar output as subtraction of a trace filtered with a rolling average of 3 seconds (**Supplementary Fig 6ciii-iv**). At the basis, these are the same procedures, linked through the Fourier transform, but details in the shape of the filtering kernels in either the real domain or the frequency domain can affect the final results of their use. For both these methods, the selection of the analysis window is crucial. Setting a too low frequency cut-off or too big rolling windows can correct photobleaching similarly to exponential decays, but maintain some artefacts (**Supplementary Fig 6civ**) and have erroneous correction at the start or end of traces (**Supplementary Fig 6ciii**). On the other hand, setting high frequency cut-off values or too small rolling windows can fully correct the baseline but might also decrease the SNR of spikes in plateau phases or mask them in the middle of noise (**Supplementary Fig 6cvi**). Moreover, because of the harsh baseline correction, most subthreshold oscillations will be lost. For all these details, choosing an appropriate baseline correction method might be the hardest task for an adequate analysis pipeline and researchers should test and adapt it for their different datasets according to their properties.

After achieving a stable baseline while retaining all signals of interest, the last analysis step is to extract specific signals such as oscillations and/or spikes. While oscillations can be analysed in a straightforward way by, for example, power spectral density analysis, spike detection can be more challenging. As spikes can have low and variable SNR, the most common way is to threshold values above a certain SNR or standard deviation (STD) and consider those as spikes (**Supplementary Fig 1g**). However, other routines based on probability or template matching have also been implemented (**Supplementary Table 1**). Choosing a suitable method will depend on the type of data collected. While thresholding might be the simplest one and is also widely used in electrophysiology analysis, it might also introduce false positive and/or false negative spike detection especially with unstable baselines or low SNR traces. In contrast, template matching might fail for low temporal resolution measurements and requires a sufficient number of spikes to create a template.

## Discussion

Voltage imaging analysis depends heavily on the specific temporal, spatial, static and dynamic features of the data. From mesoscale to subcellular spatial resolution and from slow oscillations to fast electrical activity, the required spatial-temporal resolution of the final signals leads to very distinct analysis requirements. Taking the insights gained from our analysis of different voltage imaging processing pipelines, we arrive at the conclusion that there is currently no one-size-fits-all method that can be generalizable for the analysis of all voltage imaging data. With this in mind, we present a flowchart that can help researchers to pick an optimized analysis routine for their experiments, using existing, benchmarked routines that are widely used within the community (**Fig 5**).

**Figure 5.**
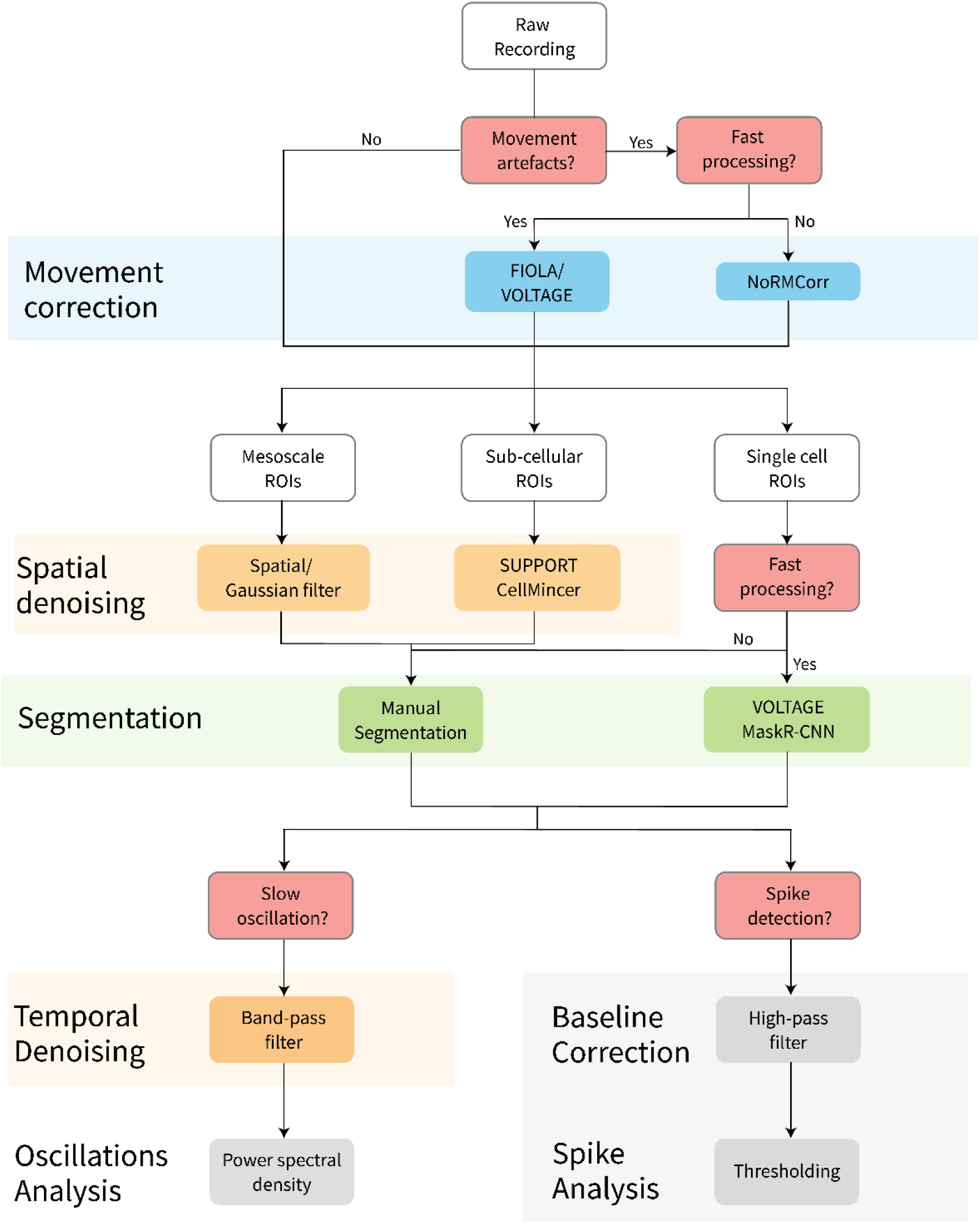
Analysis pipeline flowchart. Decision guideline depending on the different experimental settings and data types. Depending on the decision at each point, a new analysis step may be added, with recommended pipelines. White boxes represent data types. Red boxes represent decision points. Blue boxes refer to suggested motion correction methods. Yellow boxes refer to recommended denoising methods. Green boxes refer to recommended segmentation methods. Grey boxes are examples of signal processing methods.

Almost all voltage imaging data requires motion correction, especially for in vivo recordings from living animals. Movements of small areas may not heavily influence the detection of voltage fluctuations in large ROIs, cut can greatly influence signals of cells and sub-cellular compartments. NoRMCorr is the state-of-the-art routine for calcium imaging and has been also applied to voltage imaging. As voltage recordings become heavier, faster pipelines such as FIOLA or VOLTAGE can significantly improve the analyses speed, with similar results reliability than NoRMCorr. However, all motion correction pipelines will reach a limit when the movements became too large. Additionally, the movements perpendicular to the focal plane, i.e. the movements along z axis, could not be well corrected. Therefore, it is critical for researchers to first optimize experimental setups. Motion artefacts can be minimized by anesthetizing, paralyzing or restraining the animal during recordings^54, 61, 70–72^, or by decreasing the space between the coverslip of a cranial window and the animal’s brain^70^.

The following step can either be denoising or segmentation. The goal of denoising is to tackle a central challenge in voltage imaging, *i.e.* the low photon budget of the recordings. The faster the recording speed, the lower the number of photons that can be collected per frame, thus the lower the SNR of the recorded signals. For low temporal and/or spatial resolution requirements researchers can apply simple denoising methods for signal enhancement. On the other hand, for studies on cellular or subcellular ROIs focused on fast electrical activity, both the spatial and temporal resolution cannot be compromised. A solution for this challenge would be to increase the light power. However, high illumination power can lead to rapid photobleaching and to increases of tissue temperature^58^, changing the properties of the cell and possibly leading to phototoxicity. *In vivo* recordings using single-photon (1P) illumination usually require higher average power than in-vitro imaging due to the higher background fluorescence together with high illumination and fluorescence scattering. While two-photon (2P) imaging could achieve better resolution and imaging depth, it could require up to 1000x higher focal light power for the same ΔF/F as single-photon (1P)^73^.

Another way to improve cellular and sub-cellular SNR would be proper denoising of individual pixel signals. Pipelines focusing on single pixel denoising are still very limited. Three recently published methods improved denoising results in comparison to previous ones, but still offer at best modest signal improvements^38–40^, with high processing time. Denoising methods like SUPPORT and CellMincer may improve segmentation results and be particularly interesting to improve sub-cellular resolution voltage imaging data. However, with recent GEVIs developments, the fluorescence photon budget has increased sufficiently for these methods to not be needed for analysis of isolated cells. Under these circumstances, segmentation pipelines like Mask R-CNN^41^ and VOLTAGE^42^ provide accurate segmentation with as little as 1000 frames. Moreover, VOLTAGE even improves its segmentation accuracy with increasing number of frames, which is advantageous for recordings with low cell to background contrast. These pipelines are suitable for detecting individual cells, but for other ROIs, such as those of mesoscale and sub-cellular studies, manual annotations are still the most accurate segmentation routine, since each study requires analysis of different types of ROIs. Mask R-CNN and VOLTAGE are also not proven to work with highly overlapping cells. However, recent studies have developed methodologies to tackle this challenging problem^74^. Thus, current automated segmentation and denoising pipelines are not yet generalizable to high density expression patterns, arbitrary ROI morphologies, and low SNR recordings and require further improvements.

The final goal of voltage imaging is usually spike detection and oscillation analysis. Raw voltage traces from selected ROIs frequently have unstable baselines due to photobleaching and other artefacts. Frequency filters can be used to stabilize signals baseline, with the care for not removing frequencies of interest. Then, if a stable baseline is achieved, spikes can be detected by thresholding or template matching, similarly to most electrophysiology and calcium pipelines.

In neuroscience, an archetypical voltage imaging experiment involves imaging voltage changes of single neurons in awake behaving animals. In this setting we recommend motion correction by FIOLA, followed by VOLTAGE segmentation, baseline correction by frequency filtering and spike detection by thresholding. Moreover, for processing without GPU, we recommend NoRMCorr for motion correction, and manual annotation for segmentation. We also consider manual annotations to be the best option for ROIs other than single cells. We do not include any denoising method for the analysis of these signals, as the methodologies provide minimum improvements despite being computationally heavy. However, we do find denoising important for large ROIs, slow oscillation analysis and possibly sub-cellular compartment SNR increase. It is important to note that all these conclusions come from tests made using specific hardware and software settings, with publicly available datasets. While we are confident that this detailed and comprehensive methodology comparison might help researchers to better systematize their analyses pipelines, the actual speed and accuracy of specific pipelines should always be optimized for each individual application.

### Future directions

Voltage imaging is a forefront for researchers to develop new tools, from new indicators to new recording hardware and imaging procedures. The main goal is to increase SNR while using lower illumination power. With these, faster and more accurate pipelines can improve the analysis of such signals. As shown in the *denoising* section, machine learning approaches can enhance individual pixel signals. However, these methodologies are still far from optimal. Further developments in machine-learning-based approaches for denoising and signal enhancement could alleviate some of the analysis burdens for segmentation and signal processing and allow more detailed analysis of low amplitude signals, such as excitatory and inhibitory post synaptic potentials.

The increasing FOV and the number of cells recorded during populational imaging also creates a need for faster analysis routines, especially with the increasing interest in online or real-time analysis. To extract bone fide action potential waveforms, an imaging speed of over 500 fps is typically required depending on the cell type. Sub-millisecond real-time processing is computationally demanding. Although FIOLA has promised online analysis, this was focused on calcium imaging. Better hardware, together with faster routines may be necessary to achieve real-time voltage imaging analysis, but it is a topic of great interest for the scientific community. Such online analysis could pave the way for real time all optical electrophysiology, where researchers could optogenetically disrupt cells’ activity based on the ongoing activity of specific populations of cells. This could be further substantiated with patterned illumination, allowing researchers to optogenetically target different cells at different timepoints^60, 75, 76^. Such a setup would allow to answer questions about the role of individual cells inside of a population at unprecedent scale.

The continuous development of methodologies that expand the spatial and temporal resolution of voltage imaging recordings might ultimately lead to studies using three-dimensional, sub-cellular populational or multi-populational voltage imaging. Moreover, simplification or commercial setups might spread these techniques to an increasing number of labs. These and other experimental designs might become a reality in the next few years, bringing exciting new possibilities for a large variety of research topics. It is unlikely that a single analysis pipeline will be able to accurately process voltage imaging data across such a high diversity of experimental designs. To keep developing the field, researchers should focus not only on developing new tools, but also keep the analysis pipelines robust, testable, available and comparable to provide high-fidelity data.

## Methods

### Paper selection and methodologies extraction

To research methodologies used in recent years, we collected papers obtained from the SCOPUS search engine with the keywords “Voltage imag*” and “Voltage indic*”, between 2017 and 2024 (June at the date of the last search) for title, abstract and keywords. Note that using the ‘*’ symbol in SCOPUS will allow finding any matches with at least the same start, as an example: ‘Voltage imag*’ matches with both ‘Voltage imaging’ and ‘Voltage images’. We limited the search to the fields of ‘Neuroscience’, ‘Chemistry’, ‘Biochemistry’, ‘Genetics and Molecular Biology’ and ‘Multidisciplinary’, resulting in a total of 375 matches. From these we selected 277 based on title and abstracts. Moreover, we selected on the same search engine the preprints with the same keywords between 2023 and 2024, resulting in 52 matches, from which we selected 35 based on title and abstract. From the selected papers, we discarded 20 that were not open-access, 11 that were solely protocols, 5 pre-prints that were already published after peer-review and 9 that were solely notes or perspectives, giving a total of 265 papers for this review. This process is summarised in the schematic of **Figure 1A**.

We collected the analysis pipelines of the papers considered part of the *Biological questions* and *New Voltage Indicators* categories (for more information see the **Meta analysis** supplementary section) and summarised the methodologies in **Supplementary Table 1**. We classified the analysis methodologies used by researchers in 6 main categories: *Motion correction*, *Segmentation*, *Spatial Denoising*, *Temporal Denoising*, *Baseline correction* and lastly *Spike Detection.* We found that different studies required different analysis pipelines and some studies used more than one pipeline for the same step. Thus, the number of papers in each category may vary and the same paper may be present more than once in the same category.

### Datasets

To compare analysis pipelines we used publicly available datasets, so that our results can be tested by other researchers. The L1, TEG and HPC datasets were made available with the Volpy pipeline^41^ and obtained from the following link: https://zenodo.org/records/4515768. The HPC2 dataset was made available with the VOLTAGE pipeline^26^ and obtained from the following link https://zenodo.org/records/10020273. For all datasets, cell mask ground truths were provided together with the voltage imaging videos. The L1 dataset includes 9 videos of Layer 1 mouse cortex with cells expressing Voltron. The TEG dataset includes 3 videos of zebrafish tegmental area with cells expressing Voltron. The HPC dataset includes 12 videos of mouse hippocampus with cells expressing paQuasar3-s. The HPC2 dataset includes 13 videos of mouse hippocampus with cells expressing SomArchon. All datasets were recorded with regular single photon voltage imaging. HPC dataset was obtained from patterned illumination, while the remaining datasets were obtained with widefield illumination. These datasets provided variable types of recordings that allowed us to explore differences that analysis pipelines could have on the final data output.

### Hardware

All pipelines were tested in the workstations with the specifications below. Motion correction and denoising pipelines were tested in Workstation 1, while Segmentation pipelines were tested in Workstation 2.

**Table 1.**
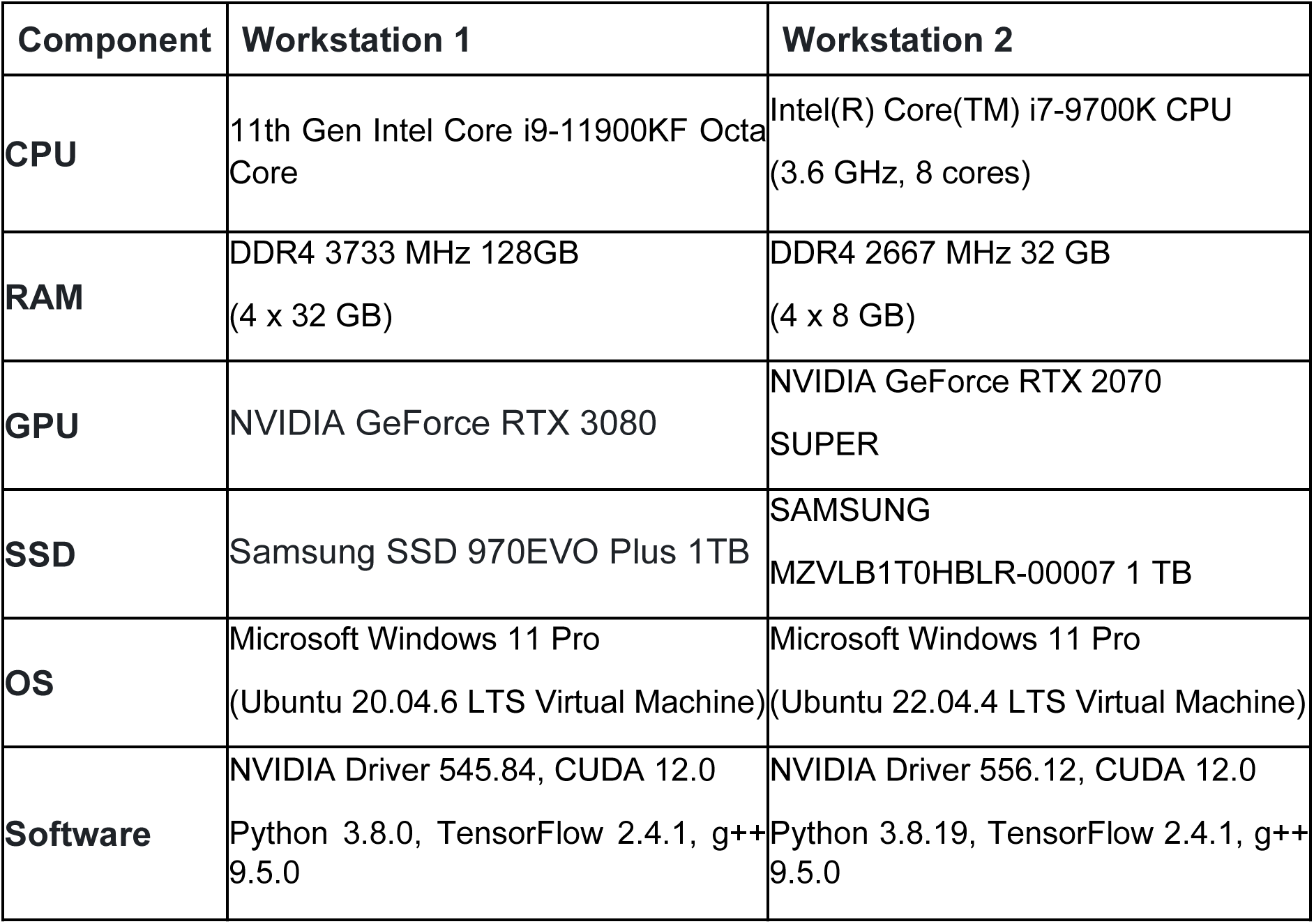
Hardware used for methodologies comparison.

| Component | Workstation 1 | Workstation 2 |
| --- | --- | --- |
| **CPU** | 11th Gen Intel Core i9-11900KF Octa Core | Intel(R) Core(TM) i7-9700K CPU (3.6 GHz, 8 cores) |
| **RAM** | DDR4 3733 MHz 128GB (4 x 32 GB) | DDR4 2667 MHz 32 GB (4 x 8 GB) |
| **GPU** | NVIDIA GeForce RTX 3080 | NVIDIA GeForce RTX 2070 SUPER |
| **SSD** | Samsung SSD 970EVO Plus 1TB | SAMSUNG MZVLB1T0HBLR-00007 1 TB |
| **OS** | Microsoft Windows 11 Pro (Ubuntu 20.04.6 LTS Virtual Machine) | Microsoft Windows 11 Pro (Ubuntu 22.04.4 LTS Virtual Machine) |
| **Software** | NVIDIA Driver 545.84, CUDA 12.0<br>Python 3.8.0, TensorFlow 2.4.1, g++ 9.5.0 | NVIDIA Driver 556.12, CUDA 12.0<br>Python 3.8.19, TensorFlow 2.4.1, g++ 9.5.0 |

### Motion correction pipelines

We tested three different pipelines, NoRMCorr, FIOLA^44^ and VOLTAGE^42^. NoRMCorr was tested inside CaImAn^48^ anaconda environment, changing solely the *max_shifts* parameter to (30,30) and the *gSig_filt* parameter to (5,5), when using this filter. We did not explore parameters since when initially tested, these parameters did not change the *max_shifts* was set to values equal or lower than (10,10) and *gSig_filt* to values equal or lower than (3,3) All other parameters were kept as default. The same values for maximum shifts were used for FIOLA. For adding the gaussian kernel, we added a convolution layer with the same gaussian kernel used for NoRMCorr, prior to the layer of shifts computation. VOLTAGE requires a Linux OS, thus we used Windows Subsystem for Linux for such tests. For this pipeline we did not change any parameter and tested it with the codes provided by the authors.

For analysis of processing time, we divided the total time of each pipeline (not counting or subtracting the timing for loading and saving results) by the number of pixels of the images processed. As the code provided by VOLTAGE authors cropped the videos in x or y coordinates, to remove black areas, we adjusted the number of pixels for these results as the number of pixels used in the motion correction processing.

### Segmentation pipelines

We evaluated two pipelines regarding fast automatic segmentation, namely Mask R-CNN^41^ and VOLTAGE^42^. We found segmentation to be heavily dependent on the parameters selected, thus for a fair comparison, we selected the parameters according to the specifications by the authors, without any further changes to the settings of either of the pipelines.

**Table 2.** Segmentation parameters used for the different datasets. The parameters in bold are the parameters that were set differently than the default settings, as suggested by the authors. For the HPC2 dataset, the frame rates (required by Mask R-CNN) were not all the same, so these values differed with different recordings: 646fps for 00_02 and 00_03 recordings; 827fps for the 01_01, 01_02 and 01_03 recordings; and 741 fps for the remaining videos.

| Method | Parameters | Datasets |  |  |  |
| --- | --- | --- | --- | --- | --- |
|  |  | L1 | TEG | HPC | HPC2 |
| VOLTAGE<br>(Bando et al., 2024) | SIGNAL_DOWNSAMPLING | 1.0 | 1.0 | **1.5** | 1.0 |
|  | SIGNAL_SCALE | 3.0 | 3.0 | 3.0 | 3.0 |
|  | SIGNAL_METHOD | **med-min** | **med-min** | max-med | max-med |
|  | TILE_SHAPE | (64, 64) | (64, 64) | (64, 64) | (64, 64) |
|  | TILE_STRIDES | **(8, 8)** | **(12, 12)** | **(4, 4)** | **(8, 16)** |
|  | TILE_MARGIN | (0, 0) | (0, 0) | (0, 0) | **(0, 0.2)** |
|  | BATCH_SIZE | 128 | 128 | 128 | 128 |
|  | NORM_CHANNEL | 1 | 1 | 1 | 1 |
|  | NORM_SHIFTS | [False, True] | [False, True] | [False, True] | **[True, True]** |
|  | PROBABILITY_THRESHOLD | **0.2** | **0.2** | **0.2** | **0.95** |
|  | AREA_THRESHOLD_MIN | **20** | **120** | **250** | **110** |
|  | AREA_THRESHOLD_MAX | **200** | **600** | **500** | **1000** |
|  | CONCAVITY_THRESHOLD | **1.3** | **1.5** | **1.3** | **1.5** |
|  | INTENSITY_THRESHOLD | **0.02** | **0.005** | **0.01** | **0.01** |
|  | ACTIVITY_THRESHOLD | **0** | **0** | **0** | **0.001** |
|  | BACKGROUND_SIGMA | 10 | 10 | 10 | 10 |
|  | BACKGROUND_EDGE | **2.0** | **3.0** | **1.0** | **1.0** |
|  | BACKGROUND_THRESHOLD | **0.003** | **0.003** | **0.003** | **0.003** |
|  | MASK_DILATION | **1** | **2** | **0** | **1** |
| Mask R-CNN<br>(Cai et al., 2024) | FRAME_RATE | **400** | **300** | **1000** | **646, 827, 741\*** |
|  | MIN_SIZE | **0** | **10** | **20** | **10** |
|  | MAX_SIZE | **1000** | **1000** | **1000** | **28** |

### Denoising pipelines

We tested three denoising pipelines, DeepVIDv2^38^, SUPPORT^39^ and CellMincer^40^, which are based on machine learning models thus requiring model training prior to testing. While SUPPORT and CellMincer only required a single video, DeepVIDv2 authors suggest using a big number of videos for model training. We used the same video of 128x512 pixels with 15000 frames from L1 dataset (L1.00.00) to train models for SUPPORT and CellMincer. For DeepVIDv2, we did not have a public dataset of the size mentioned by the authors, so we used the biggest dataset with the same dimensions we had (L1 dataset). For a fair comparison, we trained models for all three pipelines using the same dataset and, since only SUPPORT authors claimed similar performance for motion corrected and non-motion corrected videos, we used NoRMCorr motion corrected videos, both for training and testing.

We trained the models with an increasing number of iterations for each pipeline: DeepVIDv2 up to 3 iterations, SUPPORT up to 100 iterations and CellMincer up to 10000 iterations. Training times were approximately 55min per iteration for DeepVIDv2, 15min per iteration for SUPPORT and 20 seconds per iteration for CellMincer. We then denoised all datasets with the same models for each pipeline.

For computation of cells’ traces and pixel traces, we used the ground truth masks provided with the same datasets. Spikes were detected after baseline correction, using a 10Hz high pass filter, for peak amplitudes 3 times bigger than the noise standard deviation (noise level). SNR was then calculated by dividing the spikes amplitude by the noise level. The spikes detected for the cell traces were then used to calculate the SNR of individual pixel traces for all conditions.

### Parameters

*Processing speed:* For all pipelines we compared the processing speed per pixel to control for different image sizes, and video lengths.

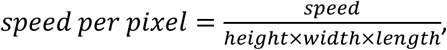

*Segmentation accuracy*: To study the accuracy of the segmentation pipelines, we computed the F1-score between the provided ground-truths and the predicted masks by VOLTAGE and Mask R-CNN for each neuron. We used the F1-score at intersection over union (IoU) of 0.3, as done in previous literature^41, 42^:

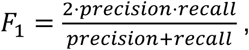

where precision and recall are calculated as:

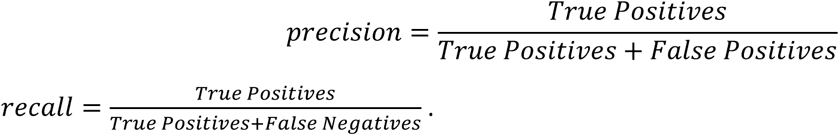

*Normalised cross correlation (NCC)*: We computed normalised cross correlation of the denoised videos by calculating the average of the first 1000 frames of each recording, normalising the frames and average image, and computing their Pearson correlation pixel by pixel.

*Baseline correction:* For all traces of the denoising comparison (**Fig 3 and Supplementary Fig3 and Supplementary Fig 4**), baselines were corrected with a 2Hz 3rd order Butterworth high-pass filter. For Signal processing tests, frequency filters were fitted also with 3rd order Butterworth filters. Rolling averages were fitted by calculating the average of the points corresponding to 0.2 or 3s surrounding a value. For the beginning and end of the traces, the rolling averages were calculated by averaging the initial or final 0.2 or 3s of the recording. Exponential and double exponential fits were computed by fitting the traces to 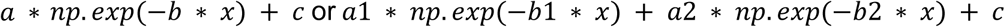, respectively.

*Spike detection*: After baseline correction we detected spikes by thresholding values higher than 3xSTD.

*SNR*: We computed SNR by first computing the baseline correction through a 2Hz high-pass filter, finding the spikes in the signals, and dividing the spike peak values by the standard deviation of the values of frames at least 10 milliseconds away from spike peaks.

### Statistical analysis

All statistical analyses, with the exception of Fig 3 and Supplementary Fig 2 (segmentation analysis), were performed by One-Way ANOVAs and Dunnet’s post-Hoc tests when One-Way ANOVAs were significant. Segmentation related statistical analysis were performed with paired samples t-tests with Bonferroni’s test for multiple comparisons. We considered data to be significantly different for p values of <0.05(*), <0.01(**), <0.005(***) and <0.001 (****).

## Author Contributions

R.S., D.B. and Z.G. conceived this study and contributed to study design. R.S. and H.B.S. conducted the analysis in the manuscript. R.S. and H.B.S. wrote the first draft of the manuscript. R.S. D.B and Z.G. revised the manuscript. D.B and Z.G. jointly supervised the project.

## Acknowledgement

This work is supported by NWO VIDI grant (Z.G., VI.Vidi.192.008) and ERC-stg grant (Z.G., 852869).

## Data availability

The data generated in the current study is available from the corresponding authors on request.

