## Supplementary Table for "Robustness and fidelity of voltage imaging analysis pipelines"

### Motion correction

| Method | Paper |
|---|---|
| NoRMCorr - rigid | (Fan et al., 2020) |
| | (Fan et al., 2023) |
| | (Wong-Campos et al., 2023) |
| | (Lowet et al., 2023) |
| | (Taxidis et al., 2023) |
| | (Brown et al., 2024) |
| | (Chen et al., 2024) |
| | (Abdelfattah et al., 2019) |
| | (Adam et al., 2019) |
| | (Kannan et al., 2022) |
| | (Evans et al., 2023) |
| | (Abdelfattah et al., 2023) |
| TurboReg | (Bando et al., 2019) |
| | (Sabater et al., 2021) |
| | (Cornejo et al., 2022) |
| | (Shiraishi et al., 2023) |
| | (Chamberland et al., 2017) |
| | (Bando et al., 2021) |
| 2D rigid fast ourier transform | (Pang et al., 2024) |
| | (Abdelfattah et al., 2019) |
| | (Platisa et al., 2023) |
| Template matching | (Piatkevich et al., 2019) |
| | (Ginebaugh et al., 2020) |
| Custom rigid transformation | (Shroff et al., 2023) |
| Trackmate software | (Bach et al., 2023) |
| Rigid registration | (Huang et al., 2024) |
| Phase correction algorithm | (Abdelfattah et al., 2019) |
| Fast variational method | (Cecchetto et al., 2021) |
| Suite2P | (Evans et al., 2023) |
| 2D particle tracing | (Shapira et al., 2021) |

### Segmentation

| Method | Paper |
|---|---|
| Manual annotation | (Ma et al., 2017) |
| | (Wienecke et al., 2018) |
| | (Tanaka & Clark, 2020) |
| | (Pan-Vazquez et al., 2020) |
| | (Milosevic et al., 2020) |
| | (Sabater et al., 2021) |
| | (Ma et al., 2021) |
| | (Kirk et al., 2021) |
| | (Walker et al., 2021) |
| | (Waiblinger et al., 2022) |
| | (Lee et al., 2022) |
| | (Platisa et al., 2022) |
| | (Böhm et al., 2022) |
| | (Costa et al., 2022) |
| | (Quicke et al., 2022) |
| | (Shroff et al., 2023) |
| | (Lowet et al., 2023) |
| | (Ma et al., 2023) |
| | (Hayward et al., 2023) |
| | (Canales et al., 2023) |
| | (Bach et al., 2023) |
| | (Campbell et al., 2023) |
| | (Morita et al., 2023) |
| | (Okray et al., 2023) |
| | (Shiraishi et al., 2023) |
| | (Shu & Jackson, 2024) |
| | (Pang et al., 2024) |
| | (Brooks et al., 2024) |
| | (Song et al., 2024) |
| | (Chamberland et al., 2017) |
| | (Kulkarni et al., 2017) |
| | (Liu et al., 2017) |
| | (Rad et al., 2018) |
| | (Kulkarni et al., 2018) |
| | (Abdelfattah et al., 2019) |
| | (Piatkevich et al., 2019) |
| | (Ortiz et al., 2019) |
| | (Abdelfattah et al., 2020) |
| | (Bando et al., 2021) |
| | (Z. Liu et al., 2022) |
| | (Evans et al., 2023) |
| | (Platisa et al., 2023) |
| | (Abdelfattah et al., 2023) |
| | (Navarro et al., 2024) |
| | (Yang et al., 2024) |
| PCA/ICA | (Fan et al., 2020) |
| | (Puppo et al., 2021) |
| | (Böhm et al., 2022) |
| | (Fan et al., 2023) |
| | (Piatkevich et al., 2019) |
| Thresholding | (Ginebaugh et al., 2020) |
| | (Alich et al., 2023) |
| | (Kulkarni et al., 2017) |
| | (Kannan et al., 2018) |
| | (Gonzalez et al., 2021) |
| | (Z. Liu et al., 2022) |
| Maximum-likelihood pixel-weighting algorithm | (Park et al., 2023) |
| | (Piatkevich et al., 2019) |
| | (Tian et al., 2023) |
| | (Ganapathy et al., 2023) |
| MaskR-CNN | (Kannan et al., 2022) |
| EXTRACT | (Kannan et al., 2022) |
| CNMF | (Kwon et al., 2017) |
| PMD-NMF | (Fan et al., 2020) |
| Watershed algorithm | (Tanaka & Clark, 2020) |
| CellPose | (Quicke et al., 2022) |
| PCA | (Wong-Campos et al., 2023) |
| K-means clustering | (Brown et al., 2024) |
| CNN | (Abdelfattah et al., 2019) |
| NMF | (Piatkevich et al., 2019) |

### Spatial denoising

| Method | Paper |
|---|---|
| Gaussian filter | (Shimaoka et al., 2017) |
| | (Rezaei et al., 2021) |
| | (Jang et al., 2021) |
| | (M. Liu et al., 2022) |
| | (Pedrosa, Nazari, et al., 2022) |
| | (Canales et al., 2023) |
| Spatial averaging | (Milosevic et al., 2020) |
| | (Jimenez-Martin et al., 2021) |
| | (Liang et al., 2021) |
| | (Zhu et al., 2022) |
| | (Milicevic et al., 2023) |
| | (Dalphin et al., 2020) |
| Spatial median filter | (Böhm et al., 2022) |
| Remove blood vessels | (Jimenez-Martin et al., 2021) |
| | (Wong-Campos et al., 2023) |
| | (Piatkevich et al., 2019) |
| PCA | (Borden et al., 2022) |
| | (Costa et al., 2022) |
| PMD | (Piatkevich et al., 2019) |
| | (Cornejo et al., 2022) |
| Bicubic interpolation | (Liang et al., 2021) |
| | (Liang et al., 2023) |
| Kalman filter | (Bando et al., 2019) |
| DeepVID | (Platisa et al., 2023) |
| Divided by low pass filter | (Evans et al., 2023) |

### Temporal denoising

| Method | Paper |
|---|---|
| Background subtraction | (Kwon et al., 2017) |
| | (Borden et al., 2017) |
| | (Hashemi et al., 2019) |
| | (Biendarra-Tiegs et al., 2020) |
| | (Lowet et al., 2023) |
| | (Bach et al., 2023) |
| | (Campbell et al., 2023) |
| | (Morita et al., 2023) |
| | (Milicevic et al., 2023) |
| | (Pang et al., 2024) |
| | (Brooks et al., 2024) |
| | (Chamberland et al., 2017) |
| | (Liu et al., 2017) |
| | (Piatkevich et al., 2019) |
| | (Ortiz et al., 2019) |
| | (Bando et al., 2021) |
| | (Shapira et al., 2021) |
| | (Gonzalez et al., 2021) |
| | (Evans et al., 2023) |
| | (Ganapathy et al., 2023) |
| | (Navarro et al., 2024) |
| Band-pass filter | (Shimaoka et al., 2017) |
| | (Fagerholm et al., 2018) |
| | (Cho et al., 2020) |
| | (Liang et al., 2021) |
| | (Rezaei et al., 2021) |
| | (Rhee et al., 2021) |
| | (Nakajima et al., 2021) |
| | (M. Liu et al., 2022) |
| | (Liang et al., 2023) |
| | (Jackson et al., 2024) |
| | (Lu et al., 2023) |
| High pass filter | (Fan et al., 2020) |
| | (Quicke et al., 2022) |
| | (Pedrosa, Song, et al., 2022) |
| | (Pedrosa, Nazari, et al., 2022) |
| | (Fan et al., 2023) |
| Low pass | (Ginebaugh et al., 2020) |
| | (Zhu et al., 2022) |
| | (Jung et al., 2023) |
| | (Lee et al., 2017) |
| Trial averaging | (Shinozaki et al., 2019) |
| | (Jang et al., 2021) |
| | (Zhu et al., 2022) |
| | (Lee et al., 2022) |
| | (Milicevic et al., 2023) |
| | (Lee et al., 2017) |
| | (Leong et al., 2021) |
| Dark Noise removal | (Pan-Vazquez et al., 2020) |
| | (Sabater et al., 2021) |
| | (Quicke et al., 2022) |
| | (Jimenez-Martin et al., 2021) |
| | (Rad et al., 2018) |
| | (Sepehri Rad et al., 2022) |
| Heartbeat filtering | (Song et al., 2018) |
| | (M. Liu et al., 2022) |
| | (Pedrosa, Song, et al., 2022) |
| | (Buchborn et al., 2023) |
| | (Meyer-Baese et al., 2024) |

### Baseline Correction

| Method | Paper |
|---|---|
| Exponential fit | (Fan et al., 2020) |
| | (Tanaka & Clark, 2020) |
| | (Milosevic et al., 2020) |
| | (Jang et al., 2021) |
| | (Sabater et al., 2021) |
| | (Rhee et al., 2021) |
| | (Platisa et al., 2022) |
| | (Zhu et al., 2022) |
| | (Cornejo et al., 2022) |
| | (Fan et al., 2023) |
| | (Wong-Campos et al., 2023) |
| | (Zhang et al., 2024) |
| | (Inagaki et al., 2017) |
| | (Abdelfattah et al., 2019) |
| | (Piatkevich et al., 2019) |
| | (Abdelfattah et al., 2020) |
| | (Alich et al., 2021) |
| | (Leong et al., 2021) |
| Double exponential fit | (Pan-Vazquez et al., 2020) |
| | (Morita et al., 2023) |
| | (Brooks et al., 2024) |
| | (Chamberland et al., 2017) |
| | (Kannan et al., 2018) |
| | (Yi et al., 2018) |
| | (Abdelfattah et al., 2019) |
| | (Leong et al., 2021) |
| | (Shapira et al., 2021) |
| | (Platisa et al., 2023) |
| | (Lu et al., 2023) |
| | (Ganapathy et al., 2023) |
| High_pass filter | (Liu et al., 2019) |
| | (Pedrosa, Nazari, et al., 2022) |
| | (Fan et al., 2023) |
| | (Shroff et al., 2023) |
| | (Lowet et al., 2023) |
| | (Huang et al., 2024) |
| | (Pedrosa et al., 2024) |
| | (Abdelfattah et al., 2019) |
| | (Shapira et al., 2021) |
| | (Kannan et al., 2022) |
| | (Z. Liu et al., 2022) |
| | (Abdelfattah et al., 2023) |
| | (Lu et al., 2023) |
| RollingMean | (Tanaka & Clark, 2020) |
| | (Pan-Vazquez et al., 2020) |
| | (Park et al., 2023) |
| | (Pedrosa et al., 2024) |
| | (Chen et al., 2024) |
| | (Jackson et al., 2024) |
| | (Huang et al., 2024) |
| | (Brooks et al., 2024) |
| | (Piatkevich et al., 2019) |
| | (Abdelfattah et al., 2023) |
| | (Platisa et al., 2023) |
| Spline | (Bando et al., 2019) |
| | (Ginebaugh et al., 2020) |
| | (Liao et al., 2024) |
| | (Piatkevich et al., 2019) |
| | (Bando et al., 2021) |
| Low pass | (Ma et al., 2017) |
| | (Milosevic et al., 2020) |
| | (Li et al., 2020) |
| | (Ma et al., 2021) |
| | (Ma et al., 2023) |
| | (Shu & Jackson, 2024) |
| | (Lu et al., 2023) |
| | (Villette et al., 2019) |
| Binomial | (Ma et al., 2021) |
| | (Canales et al., 2023) |
| | (Ma et al., 2023) |
| | (Shu & Jackson, 2024) |
| RollingMedian | (Böhm et al., 2022) |
| | (Abdelfattah et al., 2019) |
| | (Abdelfattah et al., 2023) |
| Polynomial curve | (Brown et al., 2024) |
| | (Kulkarni et al., 2017) |
| Fastsmooth (MATLAB) | (Shroff et al., 2023) |
| Low pass rectangular smoothing | (Lowet et al., 2023) |
| Assymetric Least Square Curve | (Gonzalez et al., 2021) |
| Sliding minimum | (Piatkevich et al., 2019) |
| Bottom 10th percentile | (Abdelfattah et al., 2020) |
| Savitzky-Golay | (Abdelfattah et al., 2019) |

### Spike detection

| Method | Paper |
|---|---|
| STD> 5 | (Costa et al., 2022) |
| STD>4 | (Villette et al., 2019) |
| | (Fan et al., 2020) |
| | (Shroff et al., 2023) |
| | (Fan et al., 2023) |
| | (Lowet et al., 2023) |
| | (Abdelfattah et al., 2023) |
| STD>3 | (Puppo et al., 2021) |
| | (Walker et al., 2021) |
| | (Quicke et al., 2022) |
| | (Ma et al., 2023) |
| | (Alich et al., 2023) |
| | (Kulkarni et al., 2017) |
| | (Kannan et al., 2018) |
| | (Piatkevich et al., 2019) |
| STD>3.5 | (Abdelfattah et al., 2019) |
| STD>2 | (Cornejo et al., 2022) |
| STD>1 | (Abdelfattah et al., 2023) |
| Manual threshold | (Abdelfattah et al., 2019) |
| | (Liu et al., 2017) |
| Other thresholds | (Chen et al., 2024) |
| | (Alich et al., 2021) |
| Volpy | (Taxidis et al., 2023) |
| | (Brown et al., 2024) |
| | (Kannan et al., 2022) |
| | (Han et al., 2023) |
| | (Grimm et al., 2024) |
| Kernel density method | (Huang et al., 2024) |
| | (Abdelfattah et al., 2023) |
| Spike Pursuit | (Abdelfattah et al., 2019) |
| MLSpike | (Villette et al., 2019) |
| Schmidt trigger method | (Li et al., 2020) |
| Non-linear energy operator | (Puppo et al., 2021) |
| K-means | (Walker et al., 2021) |
| Log-likelihood ratio | (Evans et al., 2023) |
| SIMA | (Liao et al., 2024) |

**Supplementary Table 1** – All papers analysed for the methodologies collected and presented in Supplementary Figure 1. References in white refer to papers related with Biological Questions. References in yellow refer to papers related with development of new voltage indicators.
