## Supplementary Figures for "Robustness and fidelity of voltage imaging analysis pipelines"

**Supplementary Fig 1**

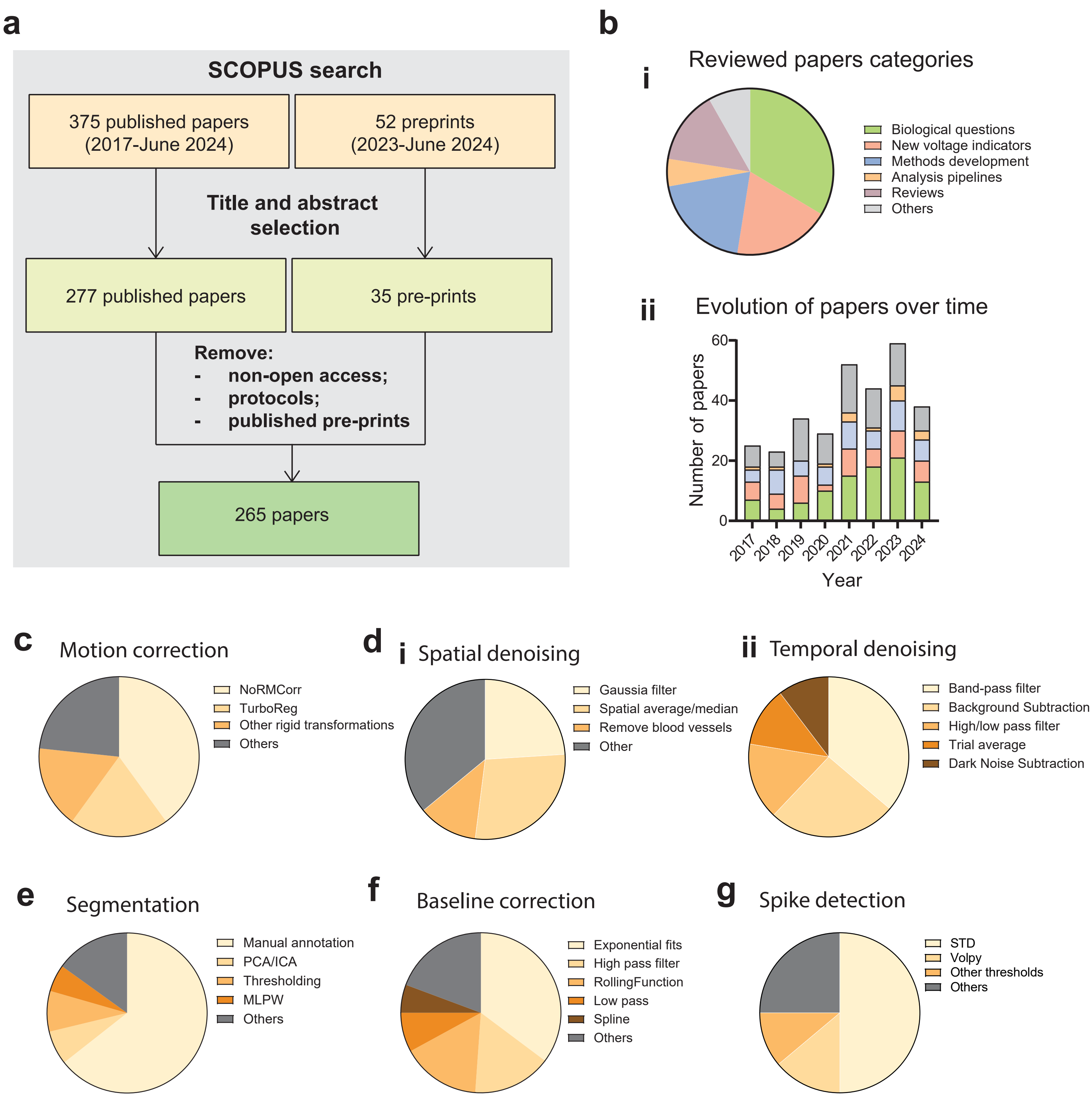

**Supplementary Fig 2**

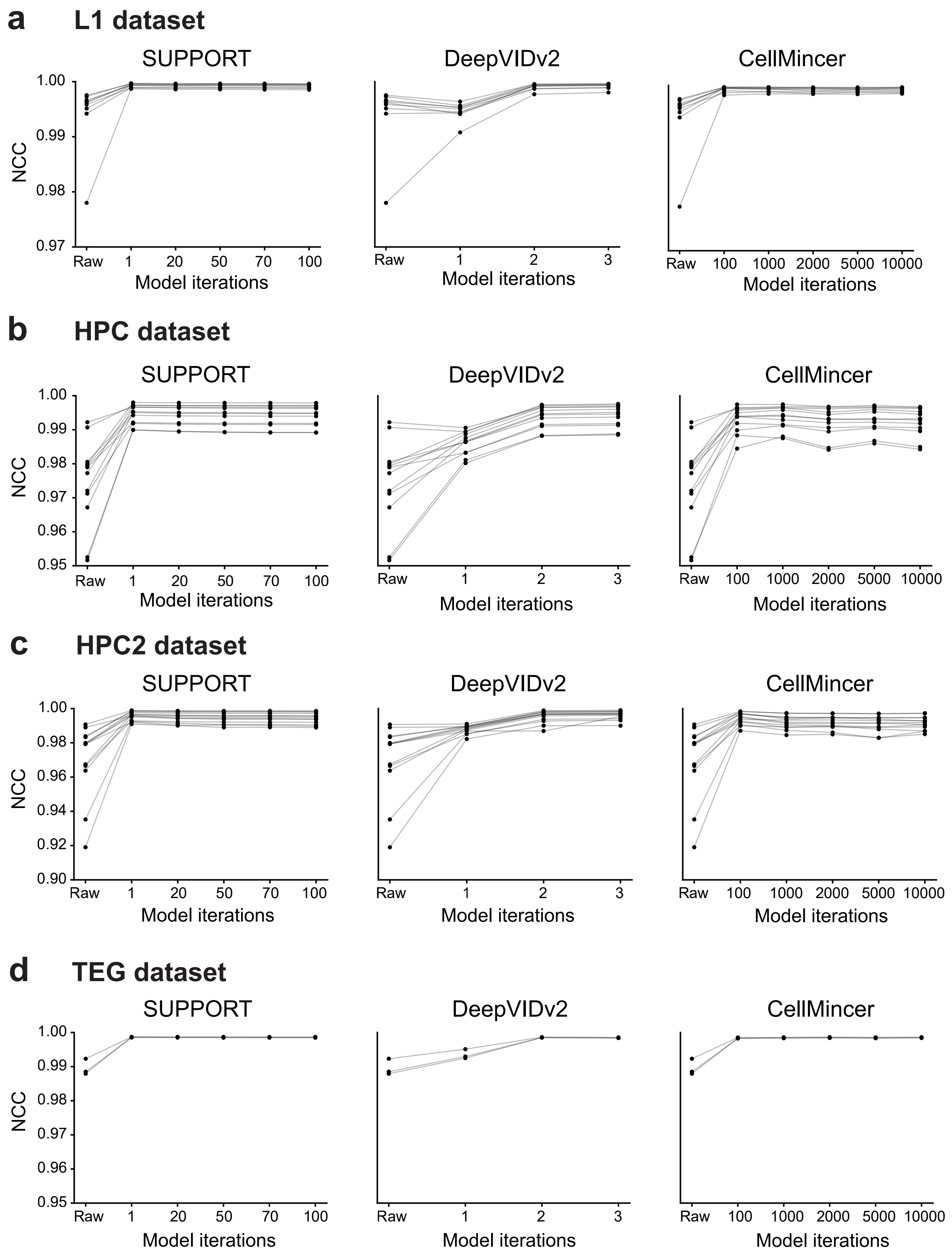

### Supplementary Fig 3

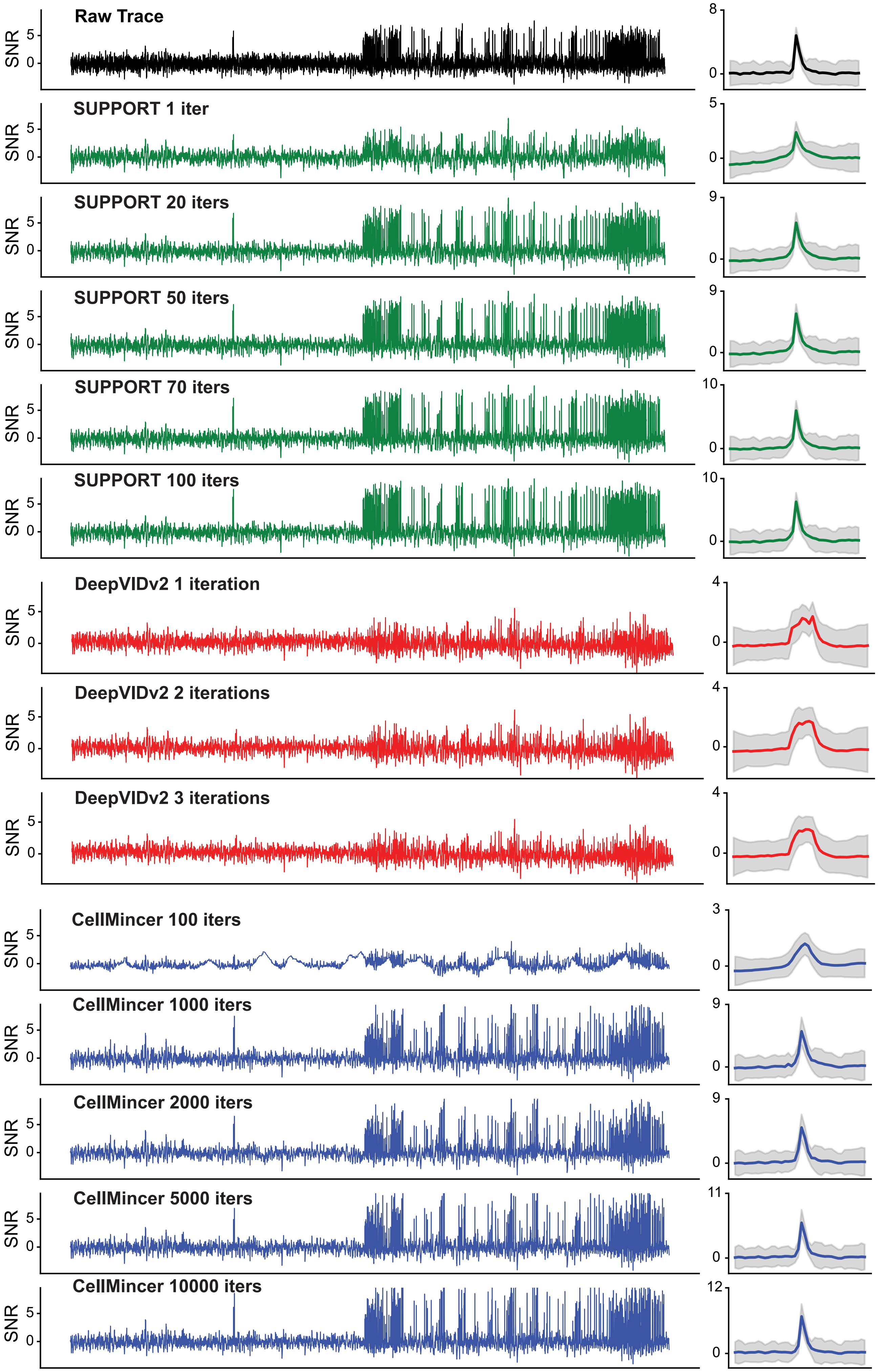

### Supplementary Fig 4

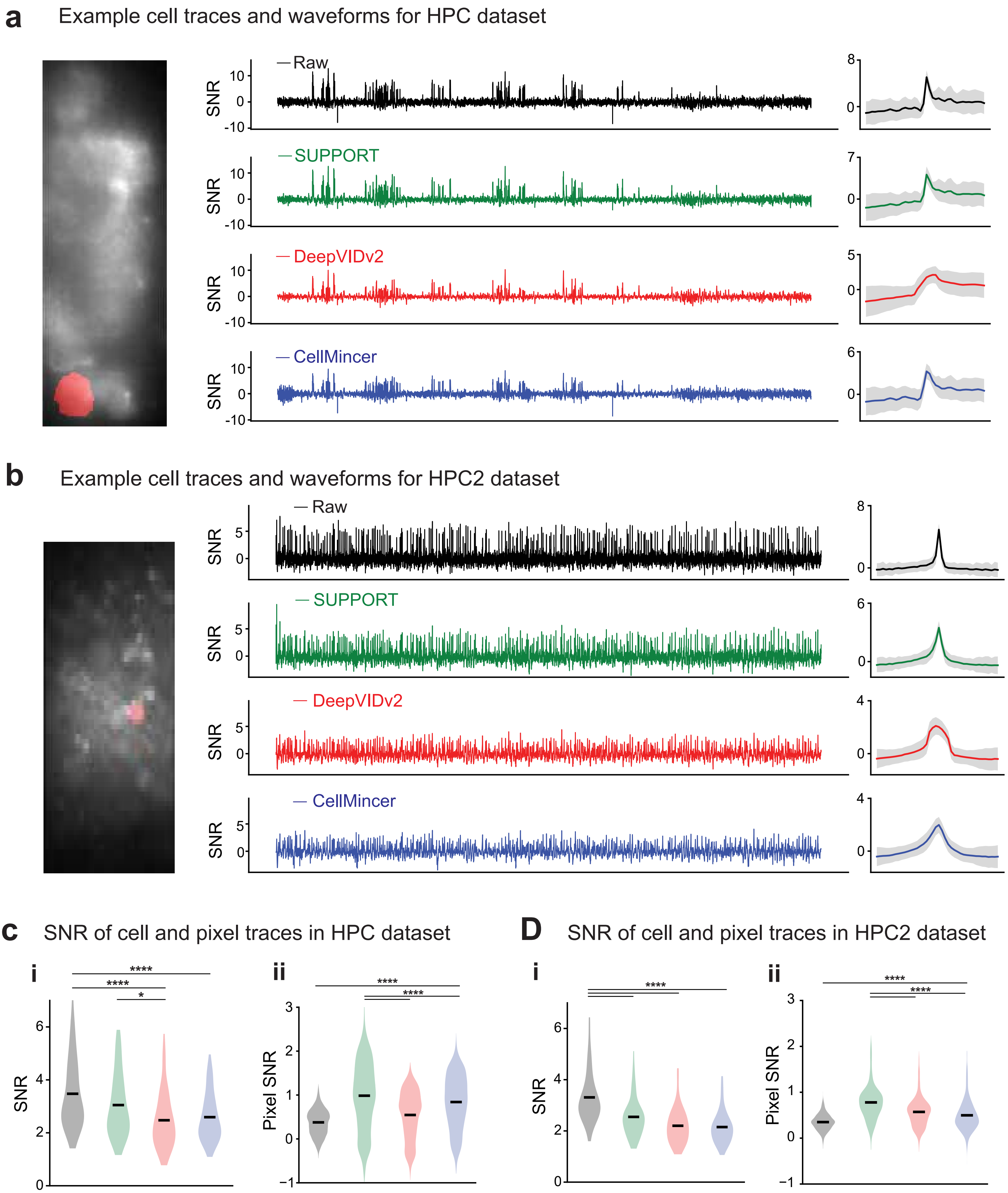

#### Supplementary Fig 5

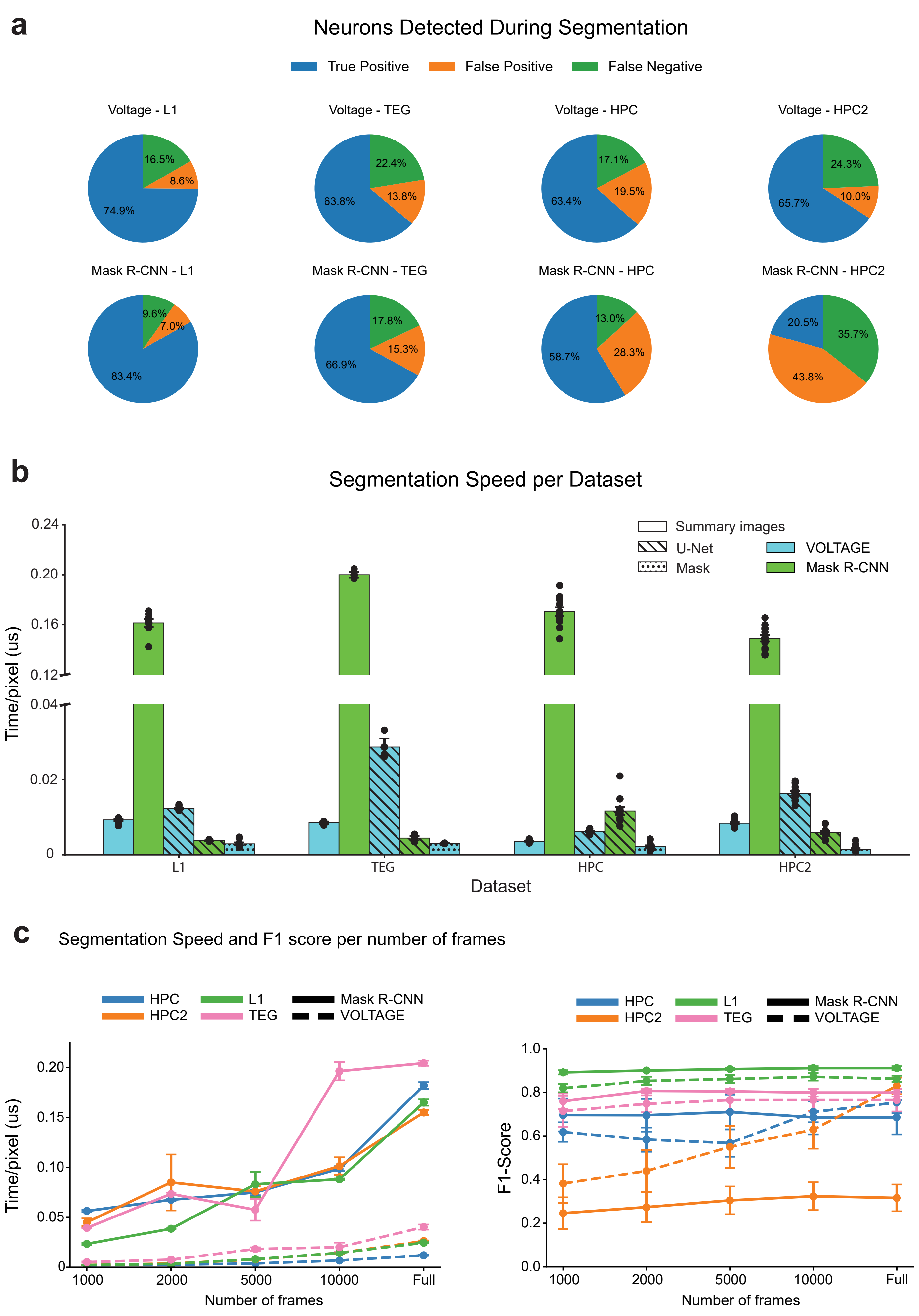

### Supplementary Fig 6

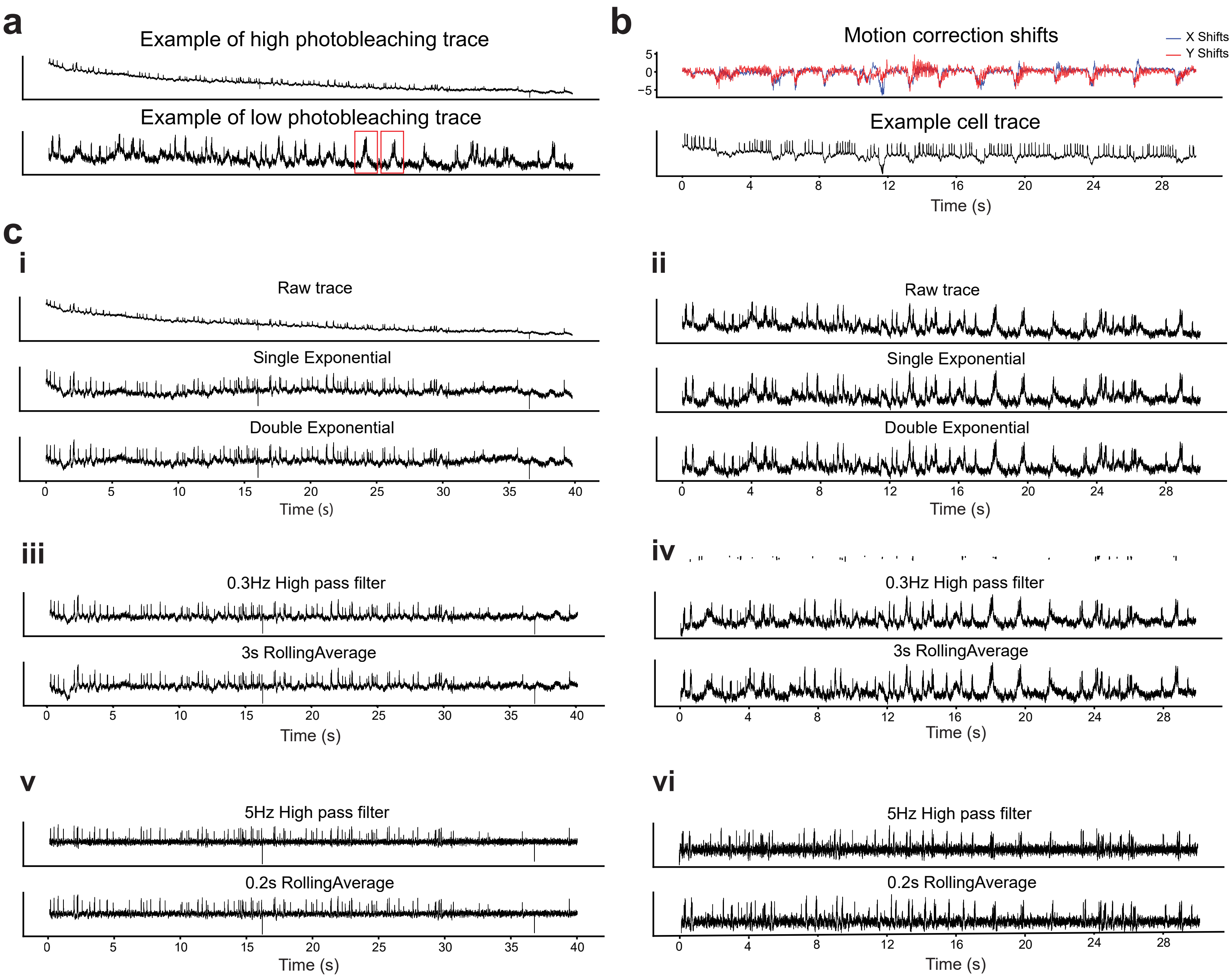

#### Supplementary Figure Legends

**Supplementary Figure 1 – A)** Paper selection method. **B)** Categorisation of papers selected according to their main topic. **i)** Ratios of papers of each category. **ii)** Evolution of papers over years, per category. **C-G)** Graphs refer only to papers from *Biological questions* and *New Voltage indicators* categories (n=147). The detailed selection is present in **Supplementary Table 1**. **C)** Ratio of motion correction pipelines used. **D)** Ratio of segmentation pipelines used. **E)** Ratio of spatial (**i**) and temporal (**ii**) denoising pipelines used. **F)** Ratio of baseline correction pipelines used. **G)** Ratio spike detection pipelines used.

**Supplementary Figure 2 -** Normalised cross correlation for each dataset of recordings denoised with SUPPORT, DeepVIDv2 and CellMincer with models trained for different number of iterations.

**Supplementary Figure 3 -** Denoised traces (left) and waveforms (right) of one example neuron with SUPPORT (green), DeepVIDv2 (red) and CellMincer (blue) with models trained for different number of iterations

**Supplementary Figure 4 -** Generalisation of denoise models for SUPPORT (green), DeepVIDv2 (red) and CellMincer (blue). **A)** and **B)** Example denoised video, cell trace and waveforms for datasets other than the one in which denoising models were trained, HPC (A) and HPC2 (B). **C) i)** Average spike SNR of all neurons in HPC dataset (n=66). **ii)** Average individual pixel SNR of all neurons in the HPC.48.08 recording of the HPC dataset (n=3800). **D) i)** Average spike SNR of all neurons in HPC2 dataset (n=63). **ii)** Average individual pixel SNR of all neurons of the 02_01 recording of the HPC2 dataset (n=1188).

**Supplementary Figure 5 - A)** Percentage of the True Positives, False Positives and False Negatives of the Segmented Neurons for each dataset. **B)** Segmentation speed per dataset (each point is one video). **C)** Segmentation speed and accuracy with increasing number of frames for all datasets.

**Supplementary Figure 6 -** Baseline Correction examples. **A)** Example of high (top) and low (bottom) photobleaching traces. **B)** Example trace with baseline fluctuations of voltage trace (bottom) correlated with movement shifts (top). **C)** Example of results of baseline correction methods in traces with high photobleaching (left) and low photobleaching (right). **i)** and **ii)** Single and Double Exponential trend filtering. **iii)** and **iv)** 0.3Hz high-pass filter and 3s rolling average trend filtering. v) and vi). 5Hz high-pass filter and 0.2s rolling average trend filtering.
